# Single nucleus RNA-sequencing defines unexpected diversity of cholinergic neuron types in the adult mouse spinal cord

**DOI:** 10.1101/2020.07.16.193292

**Authors:** Mor R. Alkaslasi, Zoe E. Piccus, Hanna Silberberg, Li Chen, Yajun Zhang, Timothy J. Petros, Claire E. Le Pichon

## Abstract

In vertebrates, motor control relies on cholinergic neurons in the spinal cord that have been extensively studied over the past hundred years, yet the full heterogeneity of these neurons and their different functional roles in the adult remain to be defined. Here, we developed a targeted single nuclear RNA sequencing approach and used it to identify an array of cholinergic interneurons, visceral and skeletal motor neurons. Our data expose markers for distinguishing these classes of cholinergic neurons and their extremely rich diversity. Specifically, visceral motor neurons, which provide autonomic control, could be divided into more than a dozen transcriptomic classes with anatomically restricted localization along the spinal cord. The complexity of the skeletal motor neurons was also reflected in our analysis with alpha, beta, and gamma subtypes clearly distinguished. In combination, our data provide a comprehensive transcriptomic description of this important population of neurons that control many aspects of physiology and movement and encompass the cellular substrates for debilitating degenerative disorders.

## Introduction

Cholinergic spinal cord neurons are essential for all aspects of motor control including voluntary contractions of the limbs and involuntary motions of internal organs. These cholinergic neurons can be divided into three main types: skeletal motor neurons, visceral motor neurons, and interneurons, with distinct functions in motor control^1^. The two types of motor neurons are particularly unusual as their cell bodies are located in the central nervous system and project to the periphery to connect the brain to the body. Skeletal motor neurons (MNs) innervate skeletal muscle to coordinate muscle contraction, drive locomotion, and mediate fine motor control. Pre-ganglionic autonomic neurons, or visceral MNs, project to autonomic ganglion neurons that in turn innervate smooth muscle and glands to control almost all physiological responses and organs of the body. The cholinergic interneurons are critical for the local spinal circuitry including regulating motor neuron excitability^2,3^. Together, these three neuronal classes comprise a relatively small population among all spinal cord cells that communicate to their target cells with the excitatory neurotransmitter acetylcholine. Previous work has shown that each of the three main classes can be divided into subtypes with specific properties and specializations^1^. However, the true diversity of spinal cholinergic neurons remains unknown.

One area of intense focus has been defining subtypes of skeletal MNs since dysfunction of this class is a major component of several diseases. Clinically, it has become clear that some subtypes are more susceptible than others to degeneration. This is particularly striking in motor neuron diseases such as spinal muscular atrophy and amyotrophic lateral sclerosis (ALS) where select MNs degenerate while others are spared^4–7^. For example, among affected neurons in ALS, skeletal MNs innervating fast-twitch extrafusal fibers degenerate earlier than those innervating slow-twitch fibers^7,8^. Most mutations linked to ALS are in widely expressed genes, yet for unknown reasons MN subtypes are more susceptible to death than other cell types, leading to the idea that cell-intrinsic characteristics determine this vulnerability^7,9^. If so, a transcriptomic definition of skeletal motor neuron subtypes could unveil potential causes of cell-type susceptibility, define better markers for studying degeneration, and provide strategies to selectively control gene expression in subsets of MNs.

With the recent advances in sequencing technologies, a few studies have transcriptionally profiled individual spinal cord neurons in development^10,11^ and in the adult^12^. But due to the technical approaches adopted, the rarity of cholinergic neurons among all spinal cord cells, as well as the large size of motor neurons, only few cholinergic neurons have been successfully sequenced^11,12^. Here, we use a genetic strategy to permanently mark cholinergic nuclei in the adult mouse spinal cord and selectively enrich them for single-nucleus RNA sequencing (snRNAseq). This approach allowed us to systematically classify cholinergic neurons and generate an atlas of their transcriptional identities. Our results exposed an array of cholinergic interneuron subtypes and revealed significant but limited molecular diversity amongst skeletal MNs, identifying new targets for exploring their function. Most surprisingly, we discovered an extensive diversity of visceral motor neuron subtypes and defined their anatomic organization along the length of the spinal cord. Together, the data provide a new view of spinal cord cholinergic neuron types, identify molecular signatures for each, and provide insights into their normal physiological functions and susceptibility to disease.

### Transcriptional profiling of adult mouse cholinergic spinal neurons

Nuclear RNA sequencing has provided an important technical advance for single cell profiling from tissues such as spinal cord that are difficult to dissociate into single cells^12,13^. We selected a single nucleus RNA sequencing strategy for this reason, and to ensure the accurate and unbiased profiling of all types of motor neurons, including those of large diameter that are typically not viably captured using single cell isolation^14^. An added advantage of the single nuclei preparation compared with single cells is that it is very rapid and therefore avoids transcriptional stress responses that can occur during tissue collection and dissociation, providing a more accurate baseline representation of cell classes^11,13,14^. However, even using snRNAseq^12^ very few cholinergic neurons were identified, reflecting their sparse representation in the spinal cord.

To selectively enrich for nuclei from spinal cholinergic neurons, we engineered mice where the cholinergic lineage in the adult spinal cord was labelled using Chat-IRES-Cre. Our genetic strategy permanently marks their nuclei with a bright fluorescent protein attached to the nuclear envelope^15^ (Figure 1). Whole tissue immunolabeling and clearing revealed bright nuclei throughout the spinal cord in the expected locations for cholinergic cells – ventral horn, lateral column, and intermediate zone (Extended Data Fig. 2a; Supplementary Video 1), thus verifying our strategy. We next proceeded to isolate nuclei from fresh tissue, harvest GFP+ nuclei via fluorescence activated cell sorting (FACS) (Extended Data Fig. 1), perform snRNA-Seq (see methods), and characterize a total of 14,738 single nuclei that met standard criteria (e.g. independent transcripts, number of genes, level of mitochondrial transcripts). Preliminary analysis revealed that the cholinergic lineage segregated into 31 distinct clusters (Extended Data Fig. 2a). Common cell type markers demonstrated that, while all of the nuclei were neuronal (*Snap25-*expressing, Extended Data Fig. 2c), 13 of the clusters were not cholinergic, reflecting excitatory or inhibitory neurons that had expressed *Chat-IRES-Cre* during development (Extended Data Fig. 2b, c). After removal of contaminating non-cholinergic neurons, analysis of the 6,941 *Chat*-expressing nuclei identified 19 transcriptionally distinct subtypes (Fig. 2a; Supplementary Table 1), exposing considerably greater diversity for this class of neurons than had been previously suspected.

**Figure 1.**
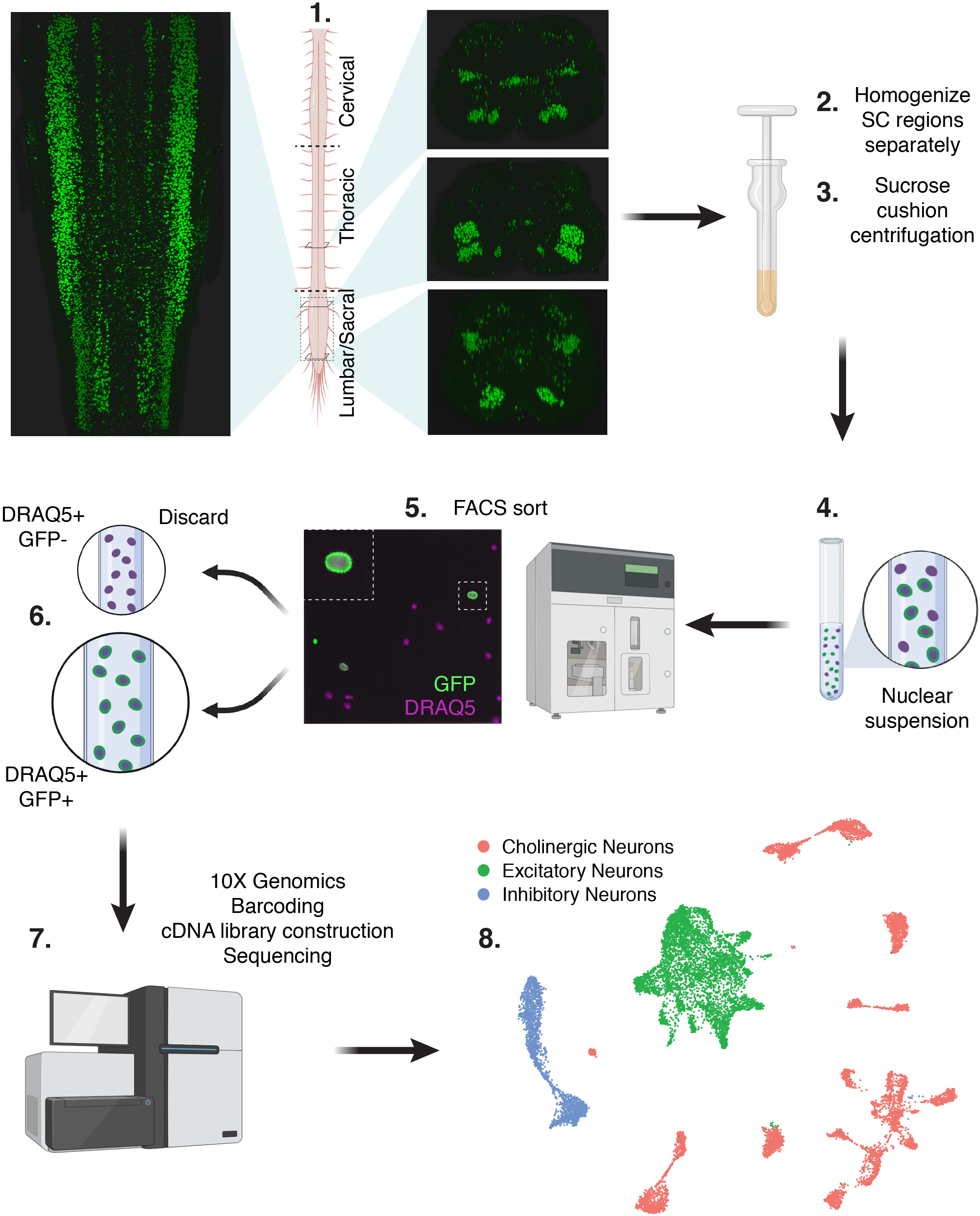
Strategy for single nucleus RNA sequencing of spinal cord cholinergic neuron. Spinal cords were extruded from Chat-IRES-Cre::CAG-Sun1-sfGFP mice and separated into 3 regions (Cervical, Thoracic, and Lumbar/Sacral) that were processed separately (1). Image from a whole cleared spinal cord immunolabeled for GFP (Extended Data Video 1). Tissue was lysed (2) and nuclei were isolated by density gradient centrifugation (3). Nuclear suspensions (4) were FACS sorted (5) to select singlet GFP-positive DRAQ5-positive nuclei (6) that were processed for single nucleus RNA sequencing using the 10X Genomics Chromium platform (7). Single nucleus cDNA libraries were sequenced together on an Illumina HiSeq (7), then analyzed (8).

**Figure 2.**
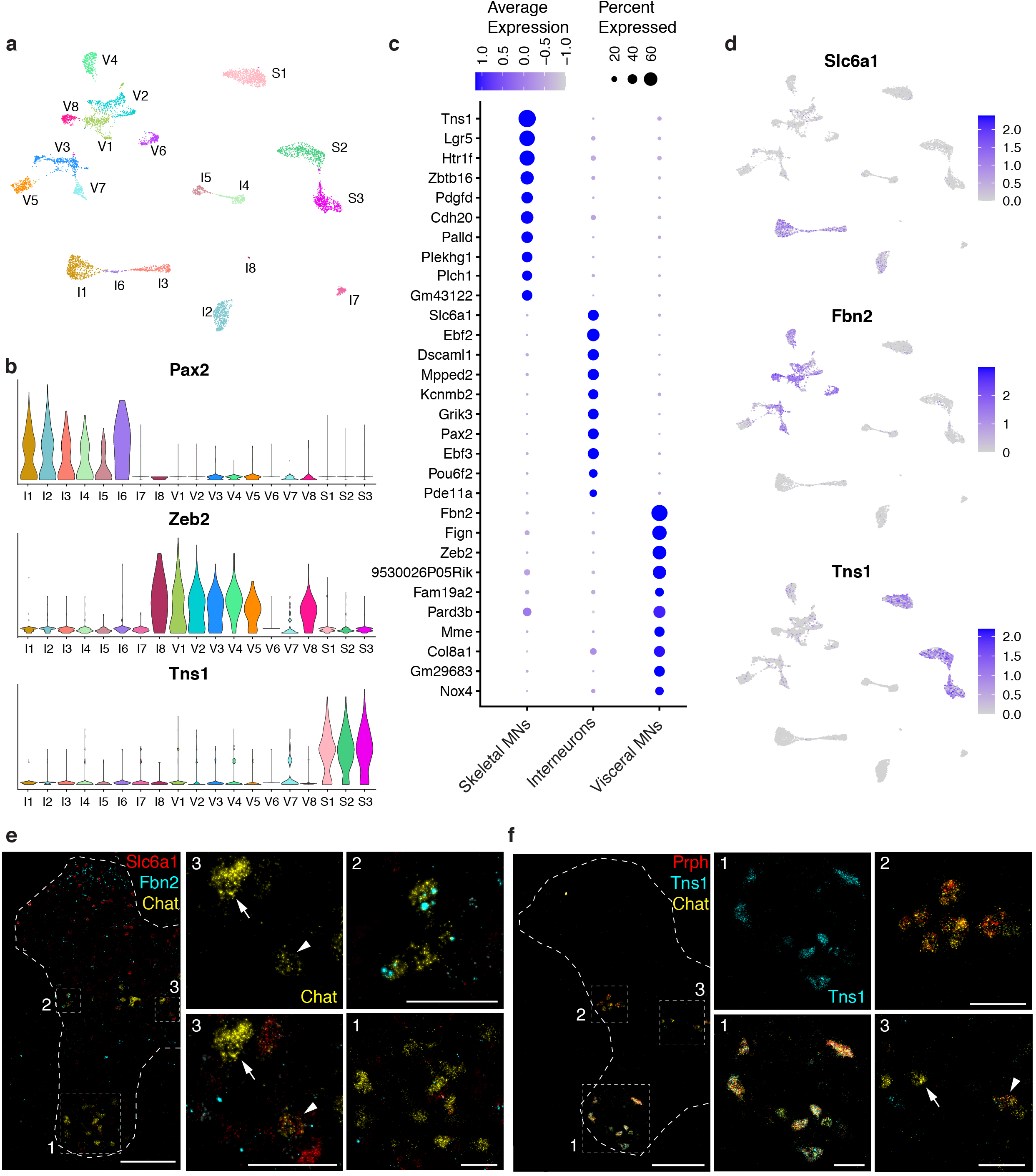
Classification of cholinergic neurons reveals the extensive diversity of motor neurons and interneurons. **a**, UMAP of all 6,941 cholinergic nuclei showing 19 distinct clusters. **b**, Expression of marker genes for cholinergic interneurons (*Pax2*), visceral MNs (*Zeb2*), and skeletal MNs (*Tns1*) in the 19 cholinergic clusters. **c**, Dot plot showing the top 10 marker genes for each main cholinergic subtype. **d**, Feature plots depicting expression of the top novel marker gene for each of cholinergic subtype. **e**, RNAScope for novel marker genes for a subset of interneurons (*Slc6a1*, red) and visceral MNs (*Fbn2*, cyan) co-expressing with *Chat* (yellow). Interneurons in the intermediate zone (box 3) are double positive for *Slc6a1* and *Chat* (arrowhead) or only express *Chat* (arrow); visceral MNs in the lateral horn (box 2) are double positive for *Fbn2* and *Chat*, and skeletal MNs in the ventral horn (box 1) are only *Chat+*. **f**, ISH for novel marker gene *Tns1* (cyan) highlights skeletal MNs, showing restricted localization to the ventral horn (box 1) and co-expression with Chat in cells with the distinctive cell body shape of skeletal MNs. Co-labeling with *Prph* (red) and *Chat* (yellow) highlights skeletal MNs (box 1) that are triple positive, visceral MNs (box 2) that are double positive for *Chat* and *Prph*, and interneurons (box 3) that are only positive for *Chat* (arrow) or occasionally double positive for *Chat* and *Prph* (arrowhead). **e, f**: thoracic spinal cord; all images are merged for the 3 colors except for *Tns1* and *Chat*, shown alone, as marked. Full single-color images and merges are shown in Extended Data Fig. 4. Low magnification image scale bars, 200 μm. High magnification scale bars, 50 μm. Boxes 1-3 are placed within ventral horn (skeletal MNs), lateral horn (visceral MNs), and intermediate zone (interneurons), respectively.

### Classification of main cholinergic neuron types

How do the 19 distinct transcriptomic classes of cholinergic neurons relate to the 3 major categories of spinal neurons? At the most basic level, the distribution of published markers for these cells in our data should help define the roles of the diverse clusters. Moreover, skeletal motor neurons, cholinergic interneurons, and visceral motor neurons each reside in distinct locations within the spinal cord (Fig. 1, Extended Data Fig. 3a, boxed areas in Fig. 2e-f;)^16–18^. Skeletal motor neurons are exclusively located in the ventral horn, with axons that exit via the ventral roots to innervate and control skeletal muscles of the body. Cholinergic interneurons, as modulators and regulators of neuronal activity, are small neurons found primarily around the central canal and the intermediate zone^2,19^. Visceral pre-ganglionic motor neurons are localized to the lateral column of the spinal cord and project to the ganglion neurons, which in turn innervate cardiac and smooth muscles for organ control^20,21^. We reasoned that, if molecular markers for each subgroup could be identified, their expression pattern in the spinal cord should be diagnostic.

We began by exploring the expression of several molecules previously assigned to subtypes of cholinergic neurons and found these markers mapped to specific clusters of neurons (Fig. 2b). For example, *Pax2*, a well-established marker for cholinergic interneurons^22^, was highly expressed in 6 clusters that we renamed I1-I6. Two small clusters, I7 and I8, expressed *Mpped2*, a gene shared with all other interneuron clusters (Fig. 2c; Extended Data Fig. 3c) but not the motor neurons. Therefore, although they were not *Pax2*-positive, we suspect I7 and I8 represent rare and unusual types of cholinergic interneurons. Meanwhile, *Zeb2*, a marker for visceral motor neurons^23^, was most highly expressed in neighboring clusters we named V1-V8. Notably, one cluster of presumptive visceral MNs based on its location in the UMAP, V6, had undetectable *Zeb2* expression, but instead shared expression of the gene *Fbn2* with the other visceral clusters, identifying this gene as a more general candidate marker for visceral MNs (Fig. 2c; Extended Data Fig. 3c). To test this prediction, we examined the expression of this gene in spinal cord sections using multiplexed fluorescent *in situ* hybridization (ISH; Fig. 2e) relative to all cholinergic neurons (*Chat*+) and most inhibitory neurons (*Slc6a1*+). Notably, *Fbn2* expression localized to a small group of *Chat*+/*Slc6a1*− neurons in the lateral column of the spinal cord where visceral MNs reside, confirming the value of *Fbn2* in identifying the full complement of visceral motor neurons.

By default, the remaining three clusters lacking *Pax2* or *Zeb2* must include skeletal MNs, for which selective markers were not known. These clusters group together in the UMAP representation and account for a significant fraction of cholinergic neurons (Extended Data Fig. 3b). Notably, the gene *Tns1* was strongly expressed in all three clusters and was essentially absent from other types of cholinergic neurons (Fig. 2b-d). ISH demonstrated that *Tns1* is robustly and selectively expressed in a group of large neurons in the ventral horn of the spinal cord, exactly where the cell bodies of skeletal MNs are located (Fig. 2f; Extended Data Fig. 4b). The *Tns1*-positive neurons invariably co-expressed two well characterized markers of cholinergic neurons, *Chat* and *Prph*^24,25^ (Fig. 2f; Extended Data Fig. 4a, b), confirming that *Tns1*-positive cells are indeed skeletal MNs. Moreover, all *Chat*-positive neurons in this area of the ventral horn co-expressed *Tns1*, thereby defining *Tns1* as a new and selective marker for skeletal MNs. In combination, these data also demonstrate that the three clusters S1-S3 represent transcriptomically distinct classes of this important type of motor neuron.

### Skeletal motor neurons

Skeletal muscle is composed of extrafusal fibers, which generate force for locomotion, and intrafusal fibers, which contain muscle spindles that detect muscle stretch and facilitate contraction. Three subtypes of skeletal MNs innervate skeletal muscle: alpha, beta, and gamma^26^. Alpha motor neurons have a large cell body diameter and exclusively innervate extrafusal muscle fibers and drive muscle contraction. Gamma motor neurons innervate intrafusal fibers, regulating muscle spindle sensitivity to stretch. Beta motor neurons are less well characterized but are known to innervate intrafusal as well as extrafusal muscle fibers and are thought to share properties with both alpha and gamma MNs^27,28^. Ample evidence shows that motor neuron disease more severely affects some subtypes of skeletal MNs^5,7^. For example, in ALS, fast-firing alpha MNs that innervate fast fatigable muscle fibers are widely described to degenerate first, both in patients and in animal models^8,29,30^, while slow-firing alpha MNs and gamma MNs are more resistant^7,31^. Discovering specific markers for each skeletal MN type and subtype would shed light on this differential vulnerability, for example by facilitating their detection and allowing their manipulation in disease models.

We hypothesized that each of the three transcriptomic clusters S1-S3 (Fig 2b, d) corresponds with one of the three functional types of skeletal motor neurons: alpha, beta, and gamma. Previous studies have defined *Rbfox3* as a selective marker for alpha MNs, and *Esrrg* and *Gfra1* for gamma MNs^32–34^. There are no known markers for beta MNs. Of the three clusters (S1, S2, S3), only cluster S2 strongly expressed *Rbfox3*, clearly assigning to it the alpha MN identity (Fig. 3a). Clusters S1 and S3 both expressed gamma markers *Esrrg* and *Gfra1*, with S1 showing slightly higher *Gfra1* expression (Fig. 3a), indicating S1 might represent gamma MNs. Because beta MNs are expected to share properties with alpha and gamma MNs^33,34^, and S3 shared transcriptional similarities with both S1 (Fig. 3a, Extended Data Fig. 5a-c) and S2 (Fig. 3b), cluster S3 most likely represents beta MNs. Several other genes reported as markers of skeletal subtypes, *Htr1d* (gammas), *Atp1a1* (alphas), and *Wnt7a* (gammas)^35–37^, did not appear restricted to any one cluster by snRNAseq (Extended Data Fig. 5a-c), perhaps reflecting the sensitivity of the single nuclear sequencing approach.

**Figure 3.**
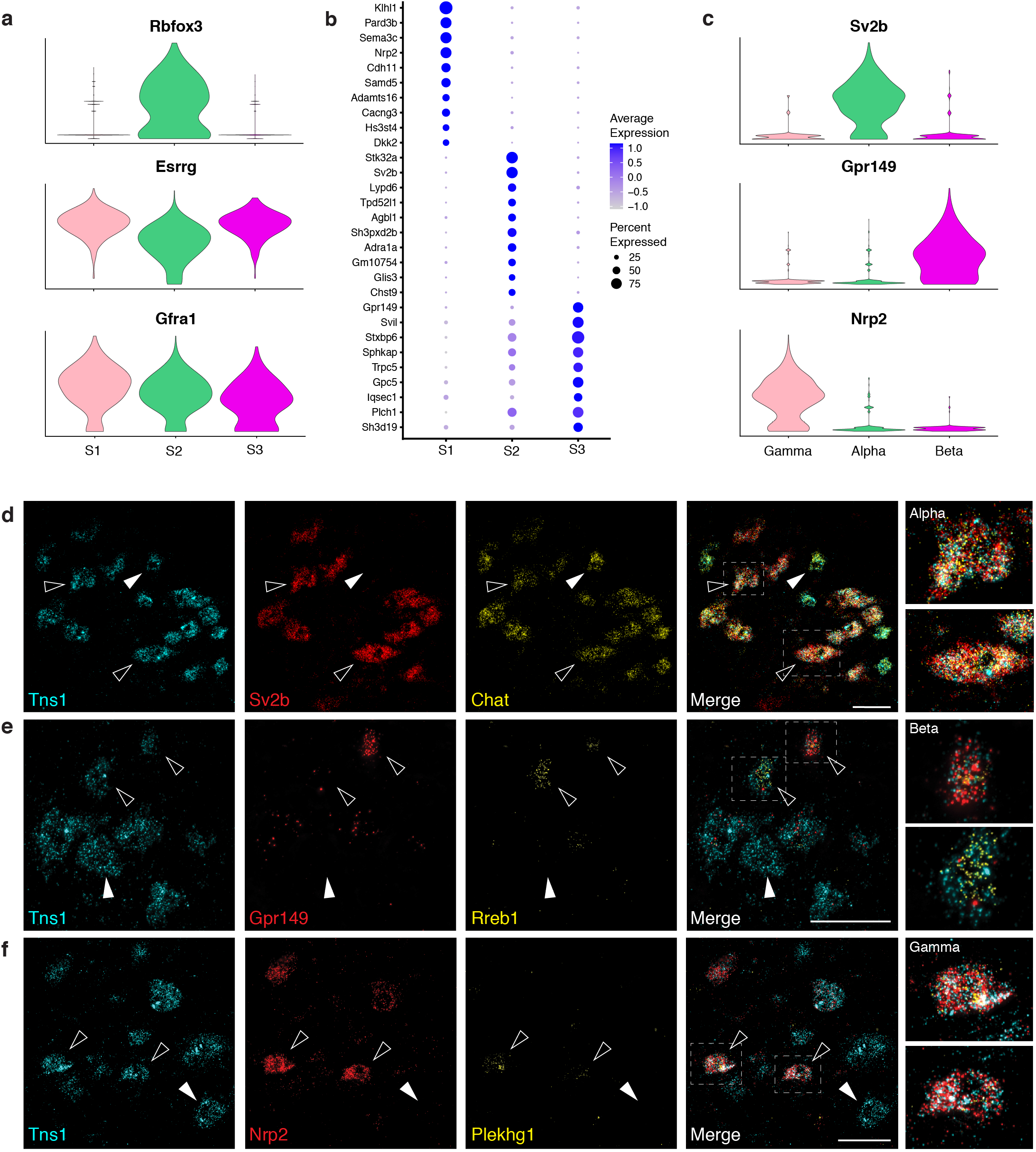
Novel markers for distinguishing alpha, beta and gamma motor neurons. **a**, Expression of known alpha and gamma markers by skeletal clusters. **b**, Dot plot for the top ten marker genes for each skeletal cluster, showing more shared expression between clusters S2 and S3. **c**, Expression of novel and specific marker genes for each skeletal MN subtype. **d**, ISH showing expression of novel alpha MN marker *Sv2b* (red) restricted to a subset of large diameter *Tns1*+ (cyan) *Chat*+ (yellow) neurons. Filled arrowhead highlights example of a *Sv2b*-negative MN of smaller diameter (i.e. beta or gamma). High magnification images of triple positive alpha MNs (open arrowheads). **e**, Smaller diameter beta MNs among *Tns1+* skeletal MNs (cyan) as revealed by expression of *Gpr149* (red) (open arrowhead, top box), or *Rreb1* (yellow; open arrowhead, bottom box) near a large diameter MN negative for both markers (filled arrowhead). **f**, Field of *Tns1*+ skeletal MNs (cyan) containing four *Nrp2*-expressing gamma MNs (red), some of which co-expressed *Plekhg1* (yellow, open arrowheads). Cells positive only for *Tns1* correspond to other skeletal MN types (e.g. filled arrowhead). ISH from ventral horn of lumbar spinal cord sections. Scale bars, 50 μm.

The expression profiles of previously identified markers support assignment of S2, S3, and S1 as alpha, beta, and gamma motor neurons, respectively, but suggest that, of these markers, only *Rbfox3* is likely to be sub-type specific. Since snRNAseq provides a rich resource for studying gene-expression profiles of different cell-types, we next used our data to identify a series of strong candidate markers for the three classes of skeletal motor neurons (Fig. 3b). For example, we predicted *Sv2b* and *Glis3* would be exclusively expressed in alpha MNs, *Gpr149* in betas, and *Nrp2* in gammas (Fig. 3b, c). Moreover, *Rreb1* would be primarily expressed in beta MNs with weaker expression in alphas, and *Plekhg1* primarily expressed in gammas with lower expression in betas (Extended Data Fig. 5f).

Multiplexed ISH revealed that these genes are expressed in MNs, thus supporting the predictions from the sequence data. Specifically, our results demonstrated that the alpha marker *Sv2b* is highly expressed in cholinergic neurons in the ventral horn (Fig 3d) and a subset of these co-expressed a second alpha-specific gene *Glis3* (Extended Data Fig. 5d). Not only do these cells express appropriate markers, including *Tns1*, but they also exhibit large diameters typical of alpha MNs. By contrast, a population of smaller diameter skeletal motor neurons expressed beta markers *Gpr149* or *Rreb1* mRNA (Fig. 3e). Similarly, we identified smaller diameter skeletal MNs expressing the gamma marker *Nrp2*, with a subset of these co-expressing *Plekhg1*, albeit at lower levels (Fig. 3f). Therefore, by multiplexing just three markers that we identified from single nuclear RNA sequence data, we were able to distinguish all the main subtypes of skeletal MNs (Extended Data Fig. 5e).

Importantly, our strategy for snRNAseq of cholinergic neurons uncovered many new genes for distinguishing between alpha, beta, and gamma MNs. The three types of skeletal MNs may exhibit different susceptibility in disease, but their specific detection has been hampered by the small number of existing markers. Even our limited survey of potential distinguishing probes (Fig. 3d-f) provides a new approach to identify all three skeletal MN types. Future localization of other genes with interesting expression patterns (Fig. 3b; Supplementary Table 2) may suggest complementary strategies and uncover even better diagnostic markers. Moreover, *Sv2b* is highly specific for alpha MNs and thus represents a significant advance.

### Sub-clustering of skeletal MN types, including distinctions between fast- and slow-firing alpha MNs

Clustering of cholinergic snRNAseq data provides strong evidence that the most prominent transcriptomic differences between skeletal MNs accounts for their division into 3 groups. However, subtypes of alpha MNs exhibit distinct electrical properties: fast-firing alpha MNs have shorter after-hyperpolarization, larger diameter axons, and more quickly adapting responses as compared to slow-firing alpha MNs^38^. In disease, certain alpha motor neurons die whereas others survive^7,39^. We next examined whether these types of distinctions were reflected in the transcriptome by re-clustering the snRNAseq data from just the skeletal MNs. This analysis distinguished 8 distinct groups that corresponded to two subclasses of alpha (A1, A2), three of beta (B1, B2, B3), and three of gamma (G1, G2, G3) MNs (Fig. 4a). The differences between subtypes within a class were more subtle (i.e. largely reflecting differences in levels of gene expression; Fig. 4b), although *Wnt7a*^37^ expression defined small and distinctive subsets of beta and gamma MNs (B3, G3) (Fig. 4a, b, Extended Data Fig. 5c).

**Figure 4.**
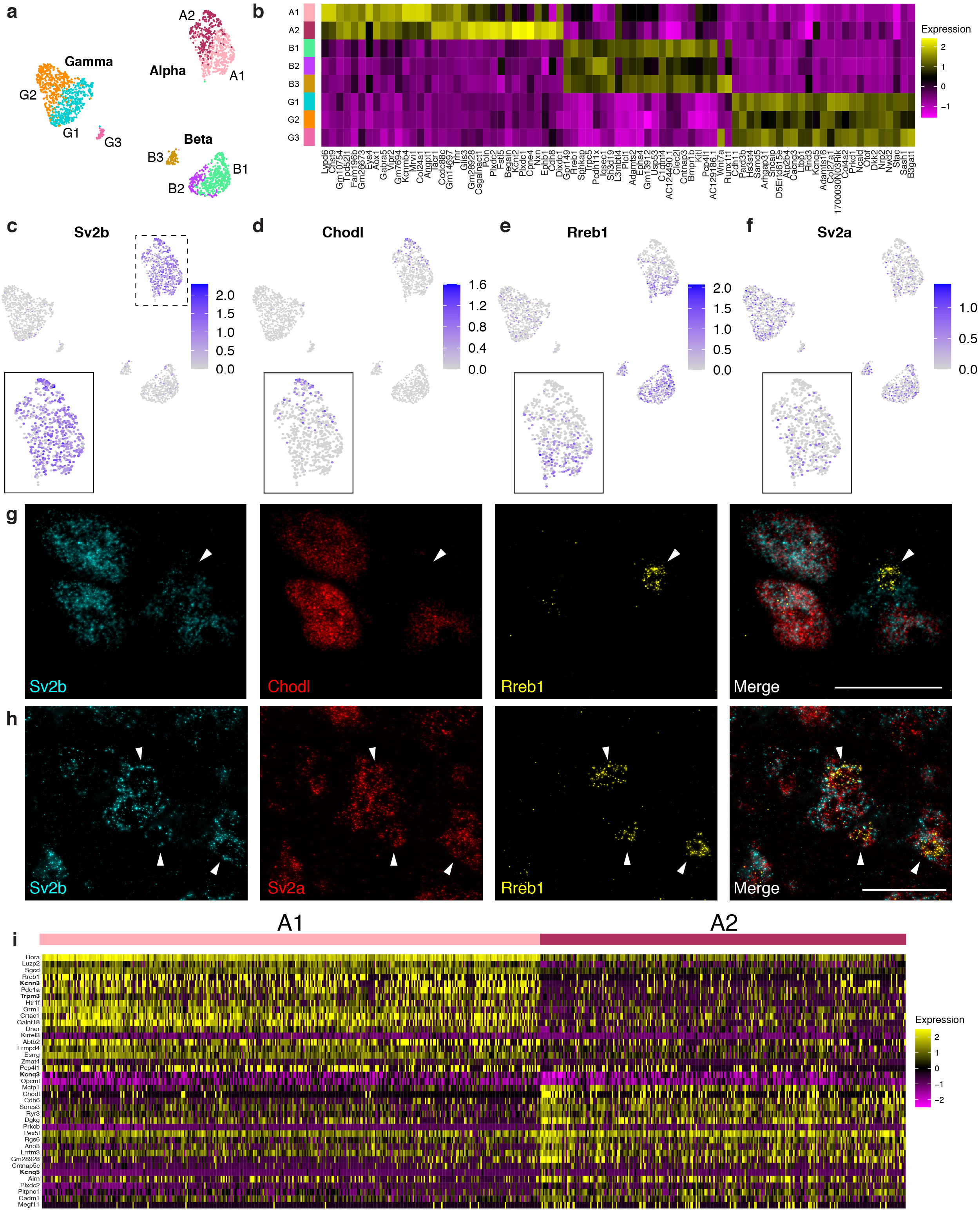
Alpha motor neurons sub-cluster into two transcriptional groups, reflecting fast- and slow-firing subtypes. **a**, UMAP of subclustered skeletal MNs by type (alpha – A1, A2; beta – B1, B2, B3; gamma – G1, G2, G3). **b**, Heatmap for expression of top genes in each skeletal sub-cluster, showing subtle transcriptional differences within each skeletal type. **c**, *Sv2b* is expressed only in the alpha cluster, spanning both sub-clusters. **d**, *Rreb1* is a marker for beta motor neurons, but also expresses in sub-cluster A1 of the alpha cluster. **e**, The known fast alpha motor neuron marker, *Chodl*, is expressed in cluster A2. Note that *Chodl* expression appears much stronger by ISH compared to its detection by snRNAseq. This is an example of how nuclear sequencing data does not always predict the robustness of an in situ signal since it does not interrogate cytoplasmic mRNA. **f**, *Sv2a*, considered a slow alpha MN marker, is broadly expressed across skeletal MNs. **g**, Cells expressing both *Sv2b* and *Chodl* depict cluster A2 alpha MNs. Arrowhead indicates a cell co-expressing *Sv2b* and *Rreb1*, representing cluster A1. Image of ventral horn in sacral spinal cord. **h**, *Sv2a* is broadly expressed in *Sv2b*+ alpha MNs. *Rreb1* labels only a subset of *Sv2b*+ alpha MNs (arrowheads). Image of ventral horn in sacral spinal cord. **i**, Heat map showing expression of top genes differentiating between *Chodl*+ and *Rreb1*+ nuclei, providing additional candidate markers for fast and slow alpha MNs. Genes in bold font are ion channels. Scale bars, 50 μm.

We focused on the subdivision of alpha motor neurons into 2 clusters (A1, A2; Fig. 4a), which we hypothesized might separate the fast- and slow-firing alpha MNs. Notably, *Chodl*, an established marker for fast alpha MNs^40^, is restricted within sub-cluster A2 (Fig. 4d). By contrast, *Sv2a*, a reported marker for slow alpha neurons^41^ appeared more prevalent within A1, although this gene was rather broadly expressed (Fig. 4f). Together, these data substantiate A1 and A2 as functionally distinct subclasses of alpha MNs and provide a new resource for understanding their biophysical properties and susceptibility in disease.

Our ISH analysis showed that *Rreb1*, a relatively selective marker for beta MNs (Extended Data Fig. 5f), was also expressed in a subset of alpha MNs (Extended Data Fig. 5e, f; Extended Data 6a). Interestingly, in the re-clustered skeletal MN analysis, *Rreb1* was primarily found in sub-cluster A1 (Fig. 4e). Moreover, ISH revealed that the *Rreb1*-positive alpha MNs all express *Sv2a* but not *Chodl* (Fig. 4g, h), suggesting that slow-firing alpha motor neurons can be identified by expression of this transcription factor. Our analysis exposed several additional candidate markers for fast- and slow-firing alpha MNs (Fig. 4i; Extended Data Fig. 6b, c). Of note, the voltage-gated potassium channel *Kcnq5* was selectively expressed in fast-firing alpha MNs (Fig. 4i; Extended Data Fig. 6d). We confirmed *Kcnq5* co-expression with *Chodl* in a subset of large diameter *Chat+* cells in the ventral horn (Extended Data Fig. 6f). Indeed, several ion channels were differentially expressed between A1 and A2 sub-clusters (*Kcnn3, Trpm3 Kcnq3, Kcnq5*; Fig. 4i; Extended Data Fig. 6b), potentially explaining their physiological differences^42,43^.

In combination, our analysis of gene expression in skeletal MNs defined markers for the three main classes, revealed that each of these can be divided into subtypes, and identified transcriptomic differences between alpha MNs that are likely to tune their firing.

### Cholinergic Interneurons

Cholinergic interneurons are found in the intermediate zone of the spinal cord where they modulate circuit function to coordinate locomotor behavior^3^, but they have not been extensively studied. The best characterized cholinergic interneurons, also called partition cells, reside near the central canal and express *Pitx2*. Elegant studies have demonstrated they are the source of cholinergic C boutons that synapse either ipsi- or contralaterally onto motor neuron cell bodies and modulate their excitability^2,3,44–46^. In our dataset, we found 8 interneuron clusters with different transcriptomic profiles and specific markers (Fig. 5a). Two, I4 and I5, express *Pitx2* (Fig. 5b), and are distinguished by the expression of *Tox* in I4 but not I5 (Fig. 5c). As predicted by the snRNAseq data, we identified examples of these two types of partition cells by ISH (Fig. 5d).

**Figure 5.**
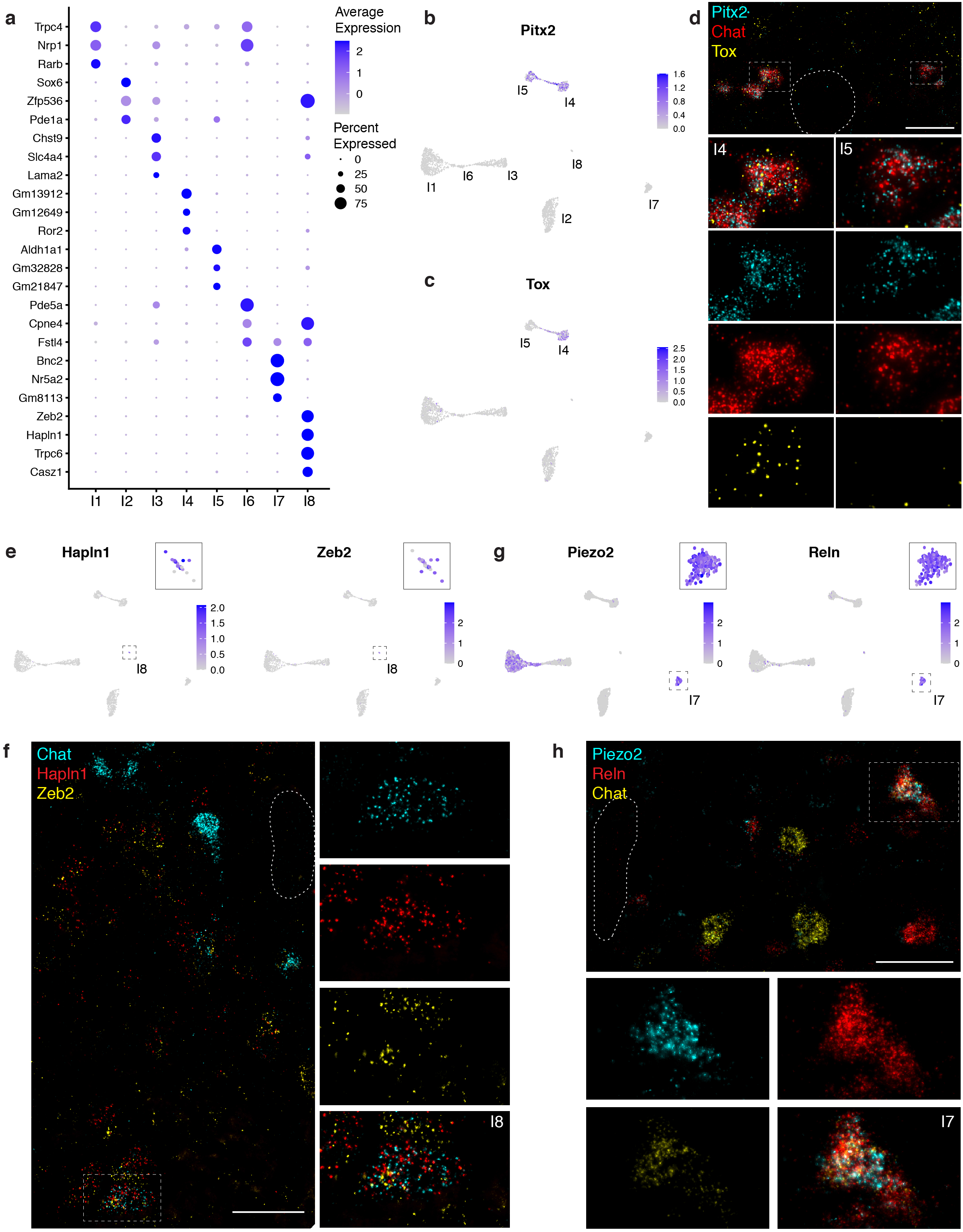
Spinal cholinergic interneurons separate into 8 transcriptionally distinct subtypes. **a**, Top markers of the 8 cholinergic interneuron clusters. **b**, *Pitx2*, a known marker for partition cells, is expressed only in clusters I4 and I5. **c**, Among interneurons, *Tox* is expressed only in cluster I4 and thus differentiates between the two *Pitx2*-expressing clusters. **d**, ISH showing expression of *Pitx2* and *Tox* near the central canal (outlined) of lumbar spinal cord. Co-expression of *Pitx2* and *Tox* in *Chat*+ cells exemplifies cluster I4 (left) whereas *Pitx2* and *Chat* without *Tox* identifies a cluster I5 cell (right). **e**, *Piezo2* and *Reln* co-expression is diagnostic of a rare population of interneurons, cluster I7. **f**, High-magnification images showing co-expression of *Piezo2*, *Reln*, and *Chat* in an I7 neuron found near the central canal (outlined) of thoracic spinal cord. **g**, Feature plots showing that co-expression of *Hapln1* and *Zeb2* in cholinergic neurons is diagnostic of cells in cluster I8. **h**, High magnification view of an example of an I8 interneuron located just ventral of the central canal (outlined) of thoracic spinal cord. Scale bars, 50 μm.

Although much more is known about partition cells, the majority of cholinergic interneurons did not express *Pitx2* (Fig. 5b). Four clusters, I1, I2, I3, and I6, accounted for almost all other cholinergic interneurons, indicating that these types of interneuron are much more diverse than was previously known. These cells were selectively labeled by *Slc6a1*, a GABA transporter (Fig 2c, d). I7 and I8 are small unusual groups of cholinergic neurons sharing expression of *Mpped2* with I1-I6 (Fig. 2c; Extended Data Fig. 3c; Extended Data Fig. 8f). To validate that these clusters represent rare types of interneuron, we defined specific combinations of markers to localize I7 and I8 (Fig 5e, g). As predicted from the sequencing, we demonstrated the existence of cholinergic neurons in the intermediate zone that co-expressed *Hapln1* and *Zeb2*, consistent with an I8 interneuron identity (Fig. 5f). ISH also revealed co-expression of diagnostic markers for I7, *Piezo2* and *Reln*, in a subset of *Chat*+ neurons (Fig. 5g). These cells localized to the intermediate zone around the central canal (Fig. 5h), substantiating their role as a new type of cholinergic interneuron.

The expression of *Piezo2* in cholinergic interneurons was surprising as this mechano-sensitive ion channel is primarily expressed in peripheral sensory neurons, where it is required for sensitivity to touch stimuli to the skin^47,48^, aspects of interoception (e.g. breathing^49^), and proprioception^48^. Interestingly, we also detected its expression in other interneuron clusters (I1, I6; Fig. 5g) as well as several visceral MN-clusters (V2, V3, V7, V8; Extended Data Fig. 8h). It will be of great interest to examine Piezo2 function in these different neuron types and to determine the types of internal mechanical stimuli that it may allow these cells to detect.

### Visceral motor neurons

The pre-ganglionic autonomic MNs, or visceral MNs, are the third main class of cholinergic cells. These neurons provide motor control in the autonomic nervous system and are responsible for relaying CNS messages that control not only involuntary movement of smooth muscles and glands but also many immediate physiological responses including heart rate, respiration, and digestion^50^. Unlike other spinal motor neurons, visceral MNs do not directly innervate their final effector organ, acting instead via ganglia in the periphery. Re-clustering visceral MNs in our dataset divided them into 14 sub-clusters (Fig. 6a) that were distinguished by select markers (Fig. 6b). Intriguingly, in addition to transcription factors, many of the markers we identified that best describe each visceral MN cluster include neuropeptides (*Ccbe1*, *Sst*, *Penk*) and genes involved with their production (*Pcsk2*). Neuropeptides modulate the function of many cells and processes, thus their differential expression in subsets of visceral MNs likely reflects the diverse functions and organs that these neurons control. Other distinguishing markers of visceral MN subtypes included secreted molecules (*Fam19a1* and *Fam163a*) and extracellular matrix proteins (*Postn, Fras1*, *Reln*, *Mamdc2*) further highlighting how their diverse gene-expression profiles are likely a reflection of their distinct physiological roles.

**Figure 6.**
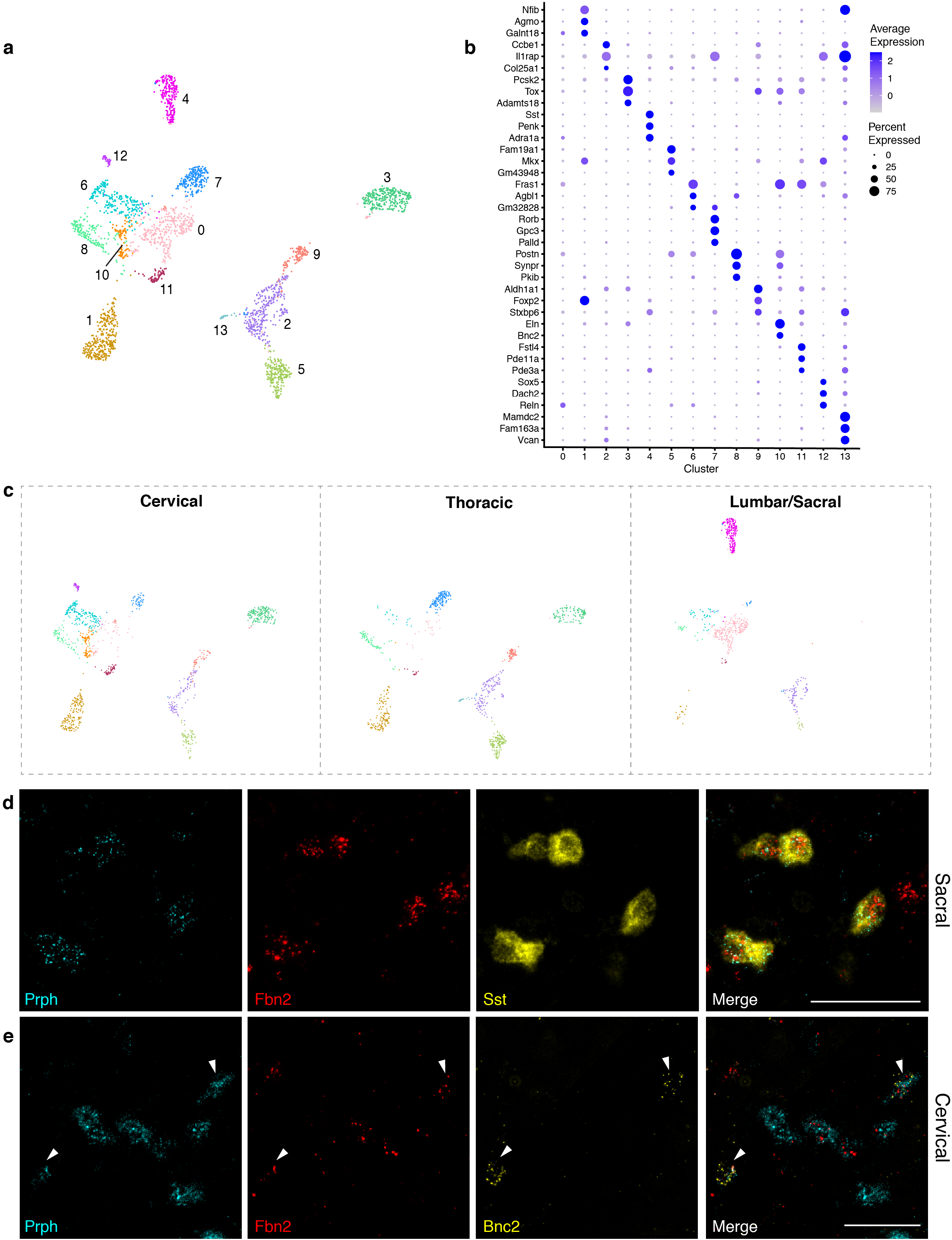
Visceral motor neurons exhibit extensive diversity in their gene expression profiles and anatomic localization. **a**, Visceral motor neurons can be re-clustered into 14 distinct sub-clusters. **b**, Dot plot showing the top three marker genes for each cluster. No markers were identified for cluster 0. **c**, UMAPs displaying the spread of visceral motor neuron clusters in each separately barcoded segment of the spinal cord. Clusters 1, 3, 5, 7, and 9 are sparse or missing from the lumbar/sacral segment; conversely, cluster 4 is unique to the lumbar/sacral segment, where cluster 0 is also most abundant. Clusters 10 and 12 are confined to cervical, cluster 6 is most abundant in cervical, and cluster 13 is confined to thoracic. **d**, ISH showing expression of *Sst*, a marker for cluster 4, restricted to lateral horn neurons in sacral spinal cord, co-expressed with *Prph* and *Fbn2*. **e**, Cluster 10 marker *Bnc2* is expressed only in cervical spinal cord in a subset of visceral MNs and co-expresses with *Prph* and *Fbn2* (arrowheads). Low magnification images showing the spinal cord region are shown in Extended Data Fig. 6. Scale bars, 50 μm.

Visceral MNs are restricted to the pre-ganglionic column in the lateral horn and thought to be primarily found at thoracic and sacral levels^1^. Moreover, immunolocalization of neuropeptides suggested that different subtypes might exist and show restricted localization along the length of the spinal cord^51^. Therefore, we designed our snRNAseq strategy to examine the distribution of cholinergic types along the length of the spinal cord by differentially barcoding nuclei from the cervical, thoracic, and lumbar/sacral cord (Fig. 1; Extended Data Fig. 3d).

Remarkably, our sequencing revealed that visceral motor neurons are abundant not only in thoracic and lumbar/sacral spinal cord but also in the cervical region (Fig. 6c), which was thought to be devoid of this type of neuron^50^. Using ISH we confirmed that diagnostic markers for visceral MNs, *Fbn2* and *Prph*, are co-expressed in the lateral column of cervical spinal cord sections (Fig 6e; Extended Data Fig. 7a). Therefore, this important class of cholinergic neurons appears more widely distributed than previously suspected. Moreover, the distribution of visceral MN clusters was strikingly diverse between cervical, thoracic and lumbar/sacral spinal cord (Extended Data Fig. 3d; Fig. 6c). Select clusters were entirely missing from certain spinal cord regions, or only present in a single region, strongly suggesting functional specialization of the neurons controlling specific organs. For example, cluster 10 marked by expression of *Eln* and *Bnc2* (Fig. 6b) was only detected in cervical derived nuclear sequence data. Importantly, ISH revealed that *Bnc2* localized to a subset of the putative visceral MNs in the cervical lateral column, further supporting our conclusion that this region of the spinal cord has a clear and specialized role in autonomic signaling.

At a more general level, our data strongly support a more complex role for visceral MNs in autonomic signaling than previously appreciated. Indeed, of 14 transcriptomically divergent clusters (Fig. 6b), 11 were unique to or more abundant in a given region (cervical: 6, 10, 12; thoracic: 13; lumbar/sacral: 0, 1, 3, 4, 5, 6, 9). Thus, their anatomic and transcriptomic specialization coincide, strongly supporting a functional role for their distinct gene expression profiles. Cluster 4 represented a large and particularly divergent group of visceral MNs that were present only in the lumbar/sacral region (Fig. 6c) and co-expressed the adrenergic receptor *Adra1a* with the neuropeptides proenkephalin (*Penk)* and somatostatin (*Sst*; Fig. 6b). Notably ISH revealed that a large subset of visceral MNs in the sacral spinal cord express the inhibitory neuropeptide *Sst* (Fig. 6d; Extended Data Fig. 7b), corresponding to pre-ganglionic neurons that have been previously described as controlling bladder and bowel function^52–54^.

## Discussion

Mammalian skeletal MNs are essential for coordinating muscle activity and all types of consciously controlled movement. They are also the cellular targets responsible for the progressive and ultimately fatal symptoms of diseases like spinal muscular atrophy and ALS. Over the past 30 years, it has become increasingly clear that diverse subtypes of skeletal MNs play distinct roles in motor control and have different susceptibility in disease. In particular, elegant developmental studies hint at considerable diversity of adult motor neurons^55^ but little is known about the true range of these cell types or the molecules defining them. Thus, to gain genetic access to skeletal MNs in animal models, the field has relied extensively on a Chat-IRES-Cre mouse that broadly targets cholinergic neurons. Here we used this line to mark cholinergic neurons and adopted a snRNAseq approach to characterize motor neuron diversity, identify new sub-types and define a wide range of genes that can be used to selectively distinguish the many different classes of adult cholinergic neurons that we discovered.

One of the most novel discoveries is the previously unappreciated number of highly distinct visceral motor neuron types. That there should exist different subtypes may not be surprising, given the diversity of organs and glands they control, ranging from the heart and lungs to the adrenal medulla, intestines, and bladder. However, it was particularly compelling to find that the different types are discretely located along the length of the cord, corresponding to a body map. An especially surprising finding was that visceral MNs extend into the cervical spinal cord, where we have clearly localized at least one subtype (*Bnc2*+ cells in the lower cervical region). All previous descriptions restrict these neurons to the thoracic and sacral regions^1^, thus it will be of great interest to further characterize these cervical visceral MNs in terms of their connectivity and function.

We also characterized a large population of cholinergic interneurons and demonstrated unanticipated diversity. Previously, only one major class of cholinergic interneurons, the *Pitx2*+ partition cells, has been studied at a functional level^2,3,45^. Our analysis divides the partition cells into two transcriptomic forms and identifies six other types of cholinergic interneurons with unknown function. Thus, the snRNAseq analysis of cells targeted by Chat-IRES-Cre mediated recombination provides a rich resource and exposes features of several unusual types of cholinergic neurons. Importantly, our data align well with a concurrent study^56^ and in combination, these data should provide the field with new approaches for selectively targeting cholinergic neuron types.

As we had hoped, our sequencing also identified skeletal MNs as three related but transcriptomically distinct groups. This clear division fits well with previous reports of select differences between alpha and gamma MNs and demonstrates that beta MNs, which have been less well characterized, form an equally abundant class. Notably, our analysis supports beta MNs as being related to both alphas and gammas^1^ but redefines them as sharing greatest similarity with a subtype of alpha MNs. The transcriptomic description of these different MN types dramatically changes the landscape of markers that can be used to distinguish them and therefore should greatly simplify their identification. Indeed, intersectional approaches using genes we validated here and *Chat*, would be extremely useful for separately targeting MN subtypes and facilitate *in vivo* dissection of their biological roles. Finally, each class of skeletal MN can be subdivided into a few transcriptomically related but distinguishable types. For the alpha motor neurons, this division appears related to their firing properties and isolates fast-firing from slow-firing subtypes. Data over the past three decades have highlighted fast-firing alpha MNs as being the most vulnerable to disease^7^, but until now it has been difficult to effectively distinguish them. We anticipate that this rich new transcriptomic dataset should help define fast-versus slow-firing MNs and expose their differential significance in health and disease.

## Methods

### Animals

Animal care and experimental procedures were performed in accordance with protocols 17-003 and 20-003 approved by the *Eunice Kennedy Shriver* National Institute of Child Health and Human Development Animal Care and Use Committee. CAG-Sun1/sfGFP mice (B6;129-*Gt(ROSA)26Sor*_*tm5(CAG-Sun1/sfGFP)Nat*_/J; Stock No: 021039^15^) were bred to the Chat-IRES-Cre::deltaNeo line (*Chat*_*tm1(cre)Lowl*_/J);Jax Stock No. 031661^57^), in which the neomycin cassette was removed to avoid ectopic expression sometimes observed in the ChAT-IRES-Cre line. This cross resulted in expression of the SUN1 fusion protein (nuclear-membrane targeting sequences of the SUN1 protein fused to 2 copies of superfolder GFP (sfGFP) followed by 6 copies of Myc) in cells that express Chat.

### Single nucleus isolation and sequencing

Spinal cords were rapidly extruded from Chat-IRES-Cre::CAG-Sun1/sfGFP mice after anesthesia with 2.5% tribromoethanol (0.5 ml/25 g body weight), decapitation and a cut through the spinal column at hip level. A PBS-filled syringe fitted with a 20G needle (7.25 mm) was inserted into the caudal end of the spinal column to flush out the spinal cord. Each cord was separated into cervical, thoracic and lumbar/sacral regions, and matching regions from 2 mice were pooled for homogenization and nuclear isolation. FACS sorted GFP+ nuclei from n=4 mice (2 females and 2 males, 8 weeks old) were pooled from each region to obtain the nuclei analyzed in this study.

The nuclei isolation protocol was adapted from Sathyamurthy et al 2018^12^. Each sample was homogenized in a Dounce Homogenizer (Kimble Chase 2 mL Tissue Grinder) containing 1 mL freshly prepared ice-cold lysis buffer (low sucrose buffer [320 mM sucrose, 10 mM HEPES-pH 8.0, 5 mM CaCl2, 3 mM Mg-acetate, 0.1 mM EDTA, 1 mM DTT] with 0.1% NP-40) applying 10 strokes with the A pestle followed by 10 strokes with the B pestle. The homogenate was filtered through a 40 μm cell strainer (FisherScientific #08-771-1), transferred to a DNA low bind 2 mL microfuge tube (Eppendorf, #022431048) and centrifuged at 300 g for 5 min at 4°C. The supernatant was removed, the pellet was gently resuspended in low sucrose buffer and centrifuged for another 5 min. The nuclei were resuspended in 500 μl 1xPBS with 1% BSA and 0.2 U/μl SUPERaseIn RNase Inhibitor (ThermoFisher, #AM2696) and loaded on top of 900 μl 1.8 M Sucrose Cushion Solution (Sigma, NUC-201). The sucrose gradient was centrifuged at 13,000 g for 45 min at 4°C. The supernatant was discarded and the nuclei were resuspended in 500 μl Pre-FACS buffer (1xPBS with 1% BSA, 0.2 U/μl SUPERaseIn RNase Inhibitor and 0.2 M sucrose). Before FACS sorting, 2.5 μl of 5 mM DRAQ5 (ThermoFisher #62251) were added.

Samples were processed on a Sony SH800 Cell Sorter with a 100 mm sorting chip and GFP+/DRAQ5+ nuclei were collected into 1.5 ml centrifuge tubes containing 10 μl of the Pre-FACS buvver. We collected ~9000, 6000, and 15000, GFP+ nuclei from the cervical, thoracic, and lumbar/sacral spinal cord samples, respectively (these nuclei were pooled from n=4 mice). Using a Chromium Single Cell 3′ Library & Gel Bead Kit v3 (10X Genomics), GFP+ nuclei were immediately loaded onto a Chromium Single Cell Processor (10X Genomics) for barcoding of RNA from single nuclei. Sequencing libraries were constructed according to the manufacturer’s instructions and resulting cDNA samples were run on an Agilent Bioanalyzer using the High Sensitivity DNA Chip as quality control and to determine cDNA concentrations. The samples were combined and run on an Illumina HiSeq2500 with Read1=98-bp, Read2=26-bp and indexRead=8. There were a total of 410 million reads passing filter. Reads were aligned and assigned to Ensembl GRm38 transcript definitions using the CellRanger v3.1.0 pipeline (10X Genomics). The transcript reference was prepared as a pre-mRNA reference as described in the Cell Ranger documentation.

### Single nucleus analysis

Sequencing data were analyzed using the R package Seurat version 3.1.4^58^ following standard procedures^59^. Outliers were identified based on number of expressed genes and mitochondrial proportions and removed from the data. Removal of outliers resulted in 14,738 total remaining cells for analysis. Each separately barcoded region of the spinal cord was separately processed by the standard methods. Briefly, the data were normalized and scaled with the SCTransform function, linear dimensional reduction was performed on scaled data, and significant principal components (PCs) were identified using the elbow method. Only significant PCs, (10-30, determined via the elbow plot method for each analysis) were used for downstream clustering. Clustering was performed using the Seurat functions FindNeighbors and FindClusters (resolution=0.6). Clusters were then visualized with t-SNE or UMAP^60^. Reference anchors were identified between each spinal cord region dataset before integration with the IntegrateData function, and integrated data were then processed by the same methods. All data was visualized with the SCT assay.

Cluster-specific marker genes were identified using the FindAllMarkers function, utilizing a negative binomial distribution (DESeq2^61^). Only positive markers were identified, and the data were down-sampled to 100 cells per cluster to facilitate comparison. Genes had to be detected in a minimum of 25% of cells and display at least a 50% log fold change. Only markers with a p-value under 0.1 were returned. In selecting top markers for each cluster, we prioritized expression in a low number of cells outside each cluster, and a higher log fold change in a large number of cells within each cluster. We filtered the markers to identify genes with expression in fewer than 30% of cells outside of each cluster and an average log fold change greater than 50% between each cluster and all other clusters and selected the genes that were least expressed outside of each cluster for visualization. In this way, we were able to select markers that are highly expressed within each cluster, while still being restricted to genes unique to each individual cluster.

### Multiplexed in situ hybridization

Spinal cord tissue was rapidly extruded as above or dissected out, immediately embedded in O.C.T. compound (Tissue-Tek), and fresh frozen on dry ice, taking care to work rapidly in order to minimize RNA degradation. The tissue was cut into cervical, thoracic, lumbar, and sacral regions that were co-embedded together for parallel processing and stored at −80°C. Blocks were sectioned into 16 μm-thick coronal slices onto positively charged slides using a Leica CM3050 S Research Cryostat. Slides were dried in the cryostat, then stored at −80°C for up to 2 weeks. Multiplexed in situ hybridization was performed according to the manufacturer’s instructions for fresh frozen sections (ACD: 320851). Briefly, sections were fixed in 4% paraformaldehyde, treated with Protease IV for 30 minutes, and hybridized for 2 hours at 40°C (HybEZ II System) with gene-specific probes to mouse *Bnc2, Chat, Chodl, Fbn2, Glis3, Kcnq5, Nrp2*, *Piezo2*, *Plekhg1*, *Prph*, *Reln*, *Rreb1*, *Slc6a1*, *Sst*, *Sv2a*, *Sv2b*, *Tox*, *Tns1*, and *Zeb2* identified from single nucleus analysis. Each probe was tested in at least n=2 mice.

Slides were imaged using a Zeiss Axiocam 506 color camera and Zeiss Apotome.2 for optical sectioning. Imaging settings were stitched using Zeiss ZEN software (blue edition). We used FIJI^62^ to generate maximum intensity projections and adjust brightness and contrast.

### Whole tissue immunolabeling and clearing by iDisco+

Whole adult spinal cords were dissected out from Chat-IRES-Cre::CAG-Sun1/sfGFP double heterozygous mice and fixed in a straightened position using 4% paraformaldehyde in PBS. To immunolabel and visualize GFP-positive nuclei within whole cleared tissue, spinal cords were processed for iDisco+ as previously described^63^. Spinal cords were immunolabeled with Chicken anti-GFP (ThermoFisher #A10262) at 1:200 during a 3-day incubation at 37°C, washed, then labeled with AlexaFluor 647-conjugated goat anti-chicken secondary antibody (ThermoFisher #A21449) at 1:200 for 2 days at 37°C. Spinal cords were embedded in 1% agarose prior to the clearing steps to facilitate their handling and imaging once cleared. Imaging was performed by light sheet microscopy using the Ultramicroscope II (Miltenyi Biotec) and the LaVision Biotec 4X objective. Images were acquired using Imspector software (single sheet, 13 dynamic focus planes, and Contrast Adaptive/Contrast algorithm). Video and 2D image renderings were generated using Arivis Vision 4D software.

### In situ probes

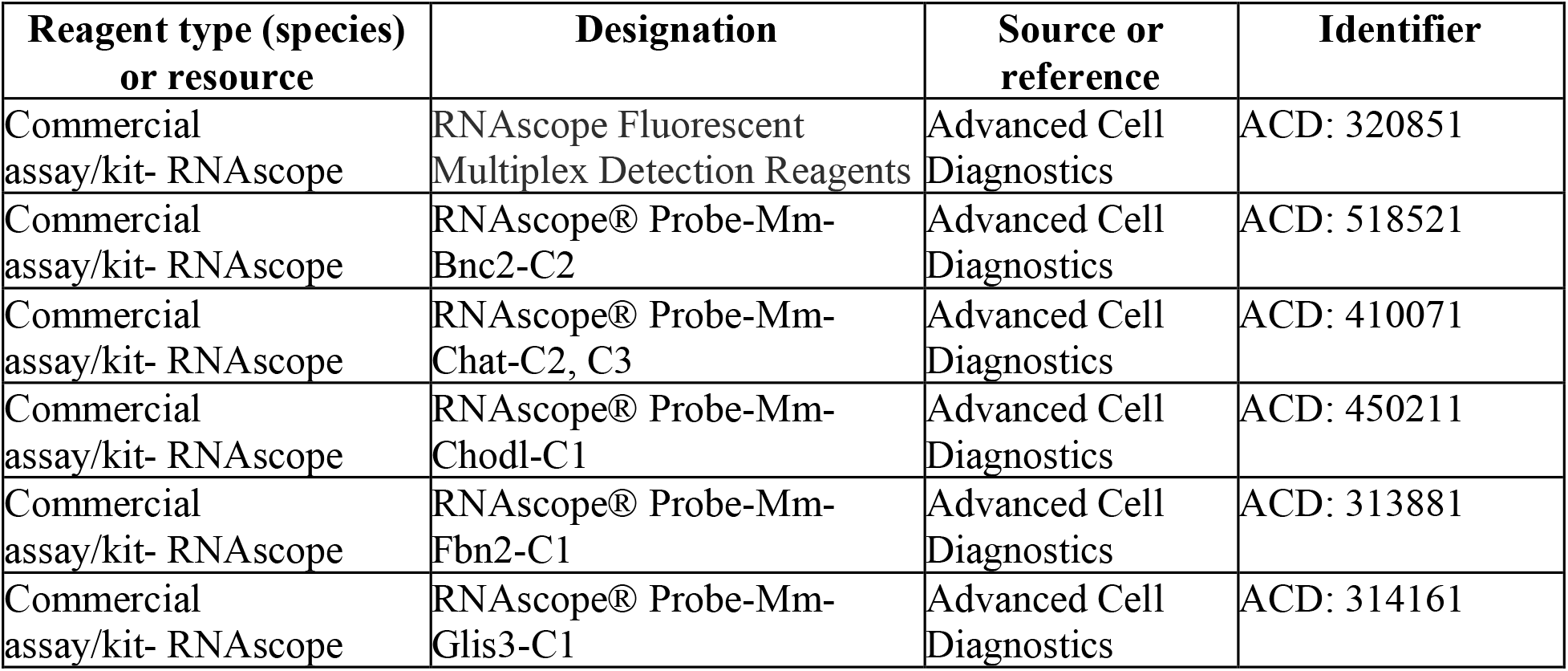
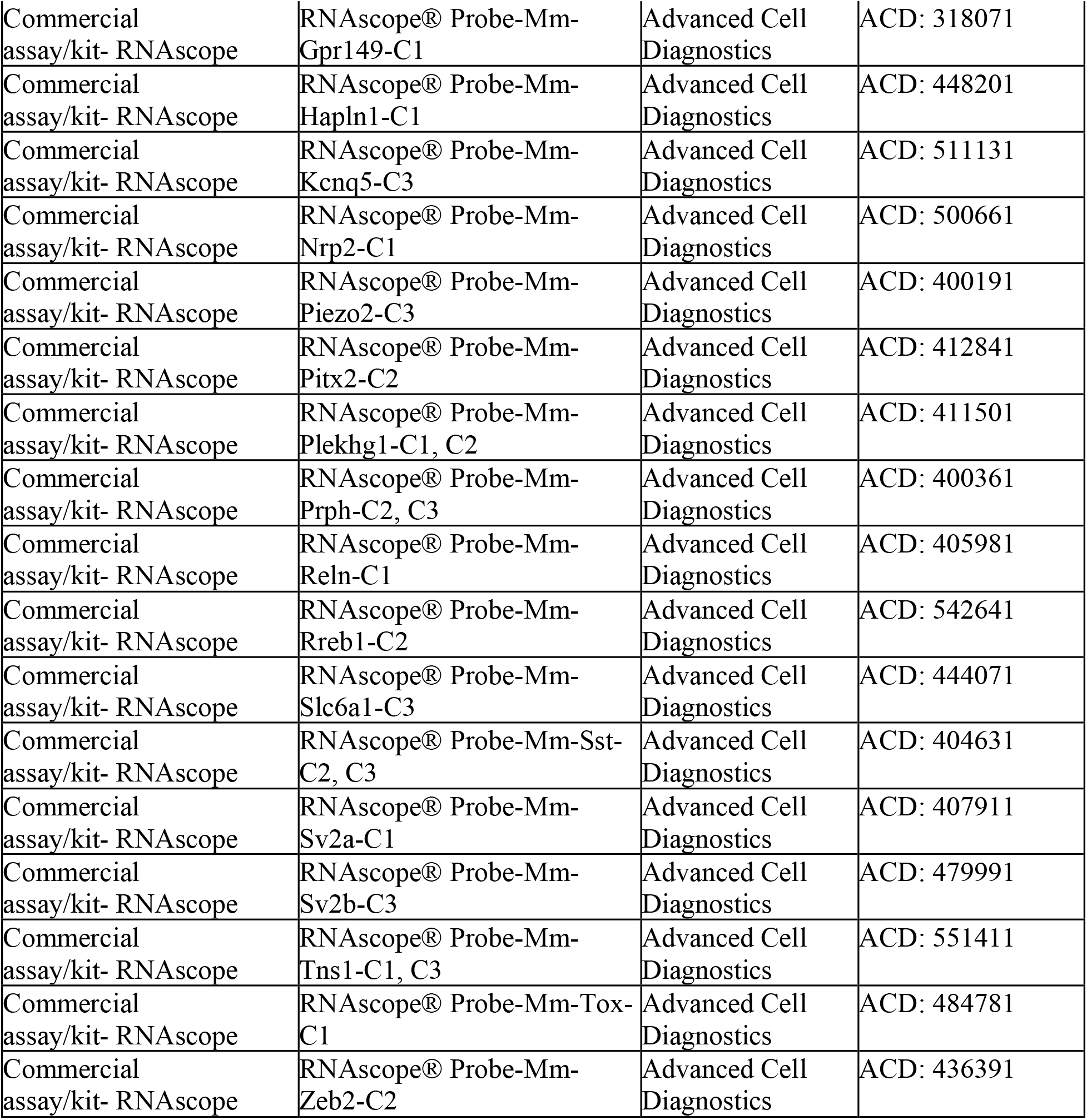

## Acknowledgements

We would like to thank intramural NIH colleagues Drs. Steven Coon, James Ibsen, and members of the Molecular Genomics Core (NICHD) for RNA sequencing and alignment to the pre-mRNA genome, Drs. Apratim Mitra and Ryan Dale of the Bioinformatics and Scientific Programming Core (NICHD) for technical advice, and Drs. Nicholas Ryba (NIDCR) and Alexander Chesler (NCCIH) for helpful discussion and comments on the manuscript.

## Author contributions

MRA, ZEP and CLP designed the experiments and wrote the paper. MRA and ZEP performed computational analysis and data validation by in situ hybridization. TJP provided advice on experimental design and expertise for sample preparation. HS generated the mouse crosses. LC and HS helped perform experiments. YZ, HS, and TJP generated the single nucleus cDNA libraries for sequencing. All authors reviewed and edited the manuscript.

## Data availability statement

The datasets generated during and/or analyzed during the current study are available from the corresponding author on reasonable request. Data will be deposited to GEO and accession codes will be available before publication.

## Code availability statement

Code used in analyzing nuclear sequencing data are found as a supplement in an R source file.

---

**Extended Data Fig. 1.**
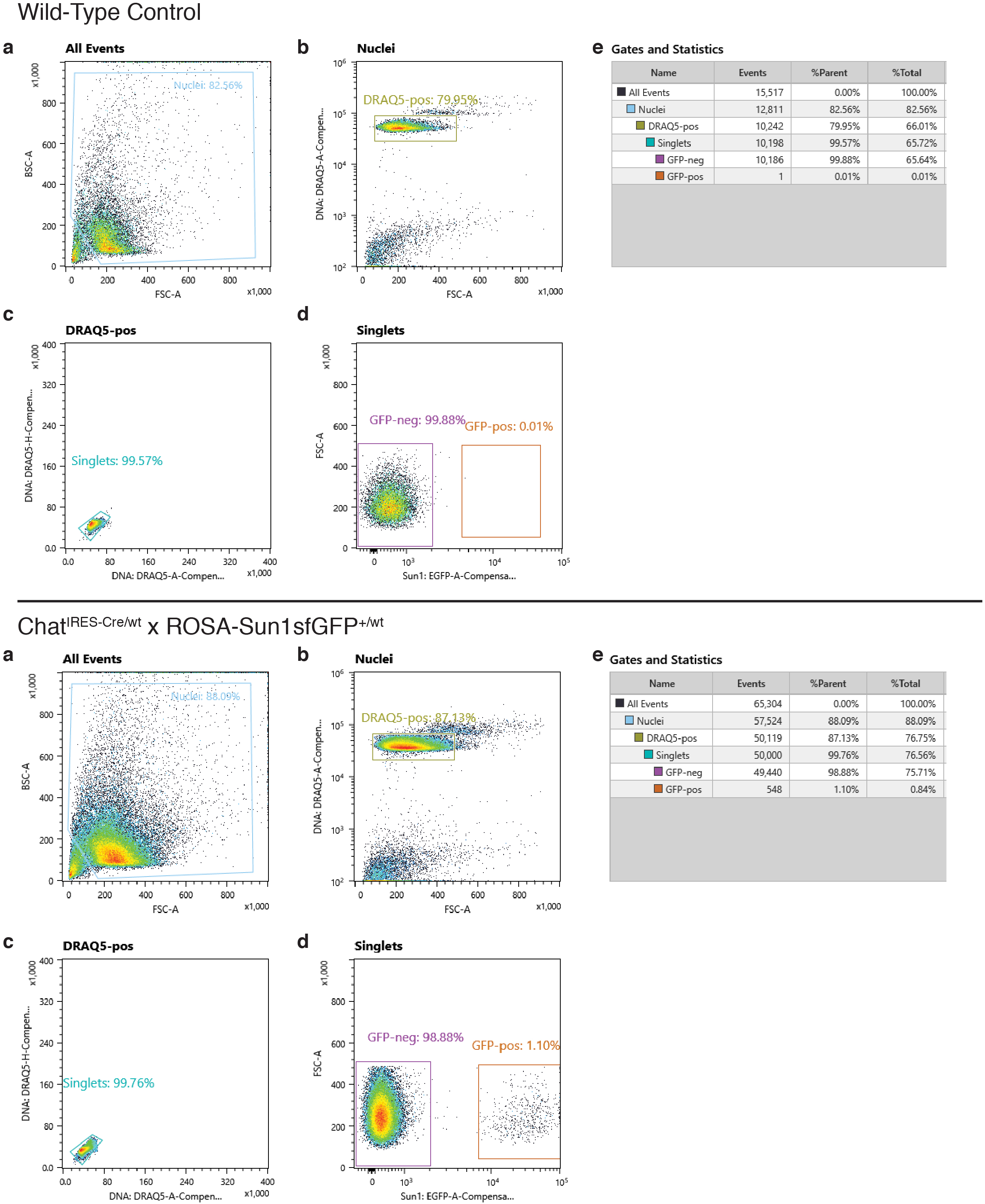
Quality control for selection of single GFP-positive nuclei by FACS sorting. Example FACS sorting summary from nuclear suspensions prepared from a wild type control sample (top), and a Chat-IRES-Cre:: CAG-Sun1sfGFP sample (bottom). **a**, Nuclear suspension visualized by FACS; **b**, gating for single nuclei by DRAQ5 positive signal; **c**, percentage of singlet nuclei among DRAQ5-positive nuclei, **d**, percentage of GFP-positive singlet nuclei; **e**, gates and statistics information.

**Extended Data Fig. 2.**
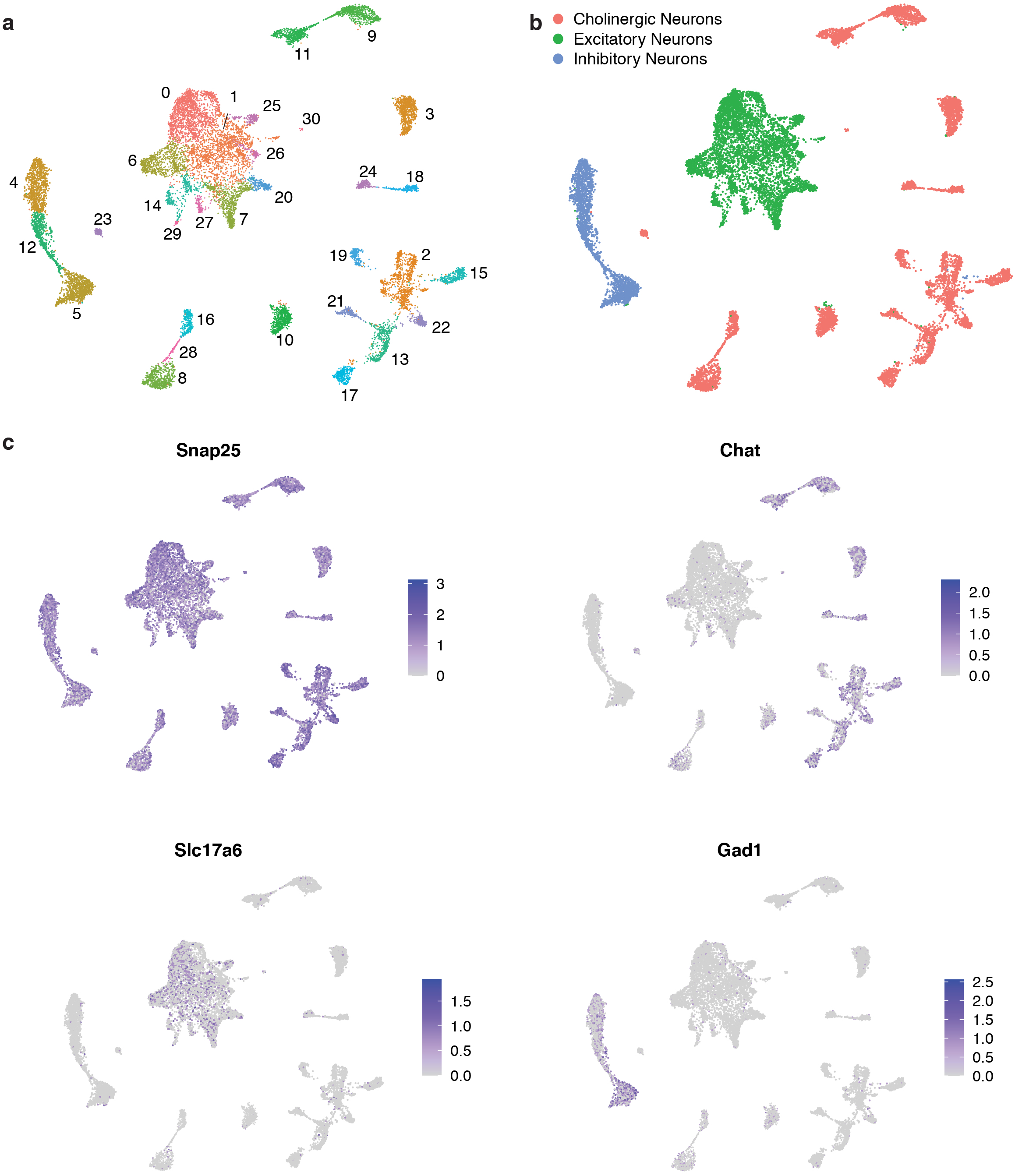
Characterization of all nuclei sequenced and assignment of cholinergic clusters. **a**, UMAP showing all nuclei analyzed, segregated into 31 distinct clusters. **b**, Clusters of inhibitory, excitatory and cholinergic neurons are color coded based on expression patters of markers shown in c. This representation is a duplication of what is shown in Fig.1 for ease of viewing. **c**, Feature plots showing expression of neuronal marker *Snap25*, cholinergic neuron marker *Chat*, inhibitory neuron marker *Gad1*, and excitatory neuron marker *Slc17a6* across all nuclei analyzed in this study.

**Extended Data Fig. 3.**
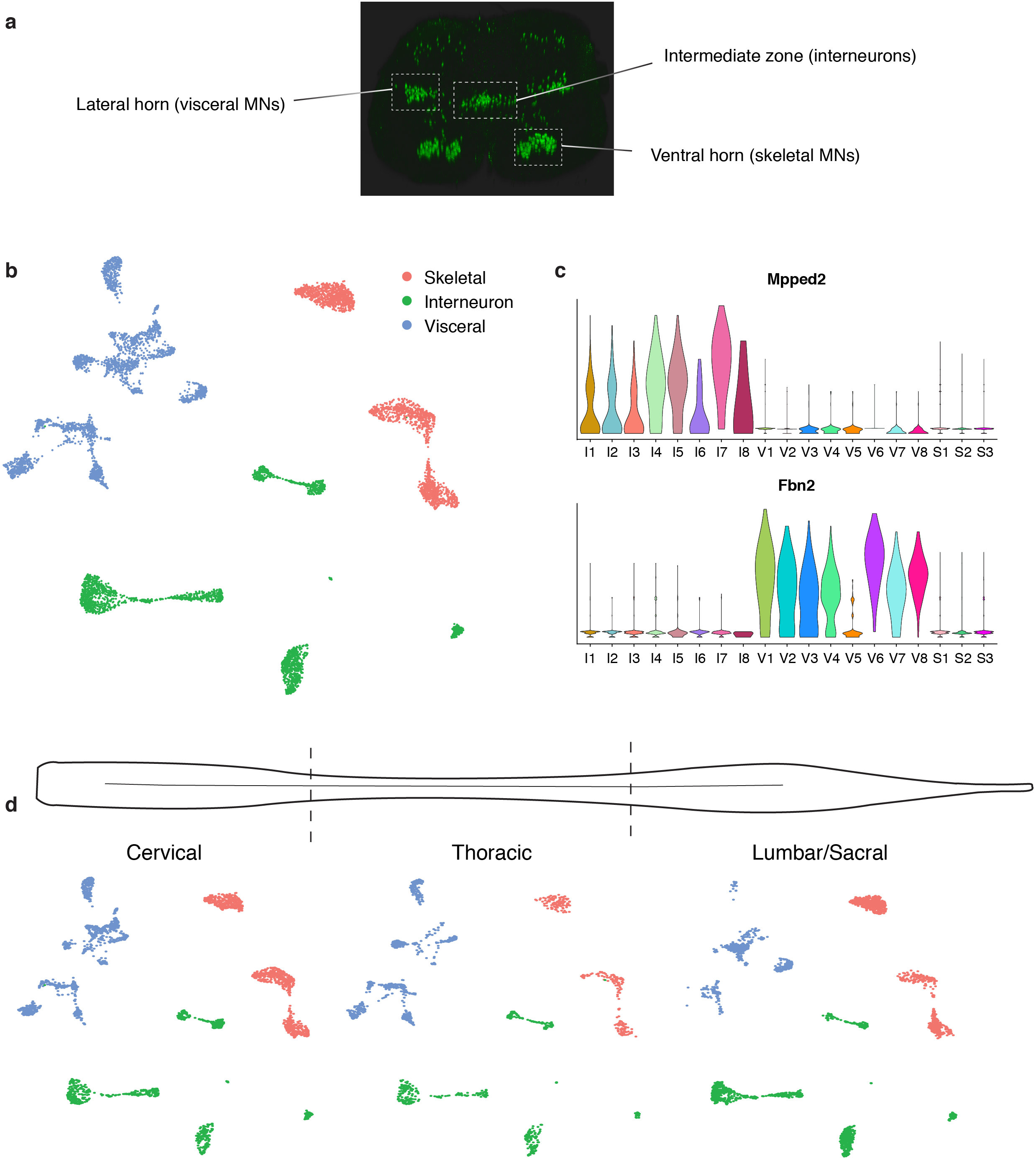
Cholinergic neuron clusters can be classified by type and vary across spinal cord regions. **a**, Lateral horn, intermediate zone, and ventral horn regions highlighted in a coronal view of thoracic spinal cord of Chat-IRES-Cre::CAG-Sun1/sfGFP mouse. **b**, UMAP showing skeletal (red), interneuron (green), and visceral (blue) clusters. **c**, *Mpped2* is a distinct marker for all cholinergic interneuron clusters, while *Fbn2* labels cluster V6 as a visceral MN cluster. **d**, UMAPs of cells detected in each spinal cord region (cervical, thoracic, and lumbar/sacral) showing variability in cholinergic neuron subtype distribution throughout the spinal cord. The visceral MNs showed striking localization differences of subtypes along the length of the cord (see Fig. 6). In contrast, interneurons and skeletal MNs showed much less variability by segment, with a similar presence of each cluster in each of the 3 regions. The only noticeable difference was that skeletal MNs were predominantly located in the cervical and lumbar regions, as to be expected due to the larger number of limb-innervating skeletal MNs in these segments.

**Extended Data Fig. 4.**
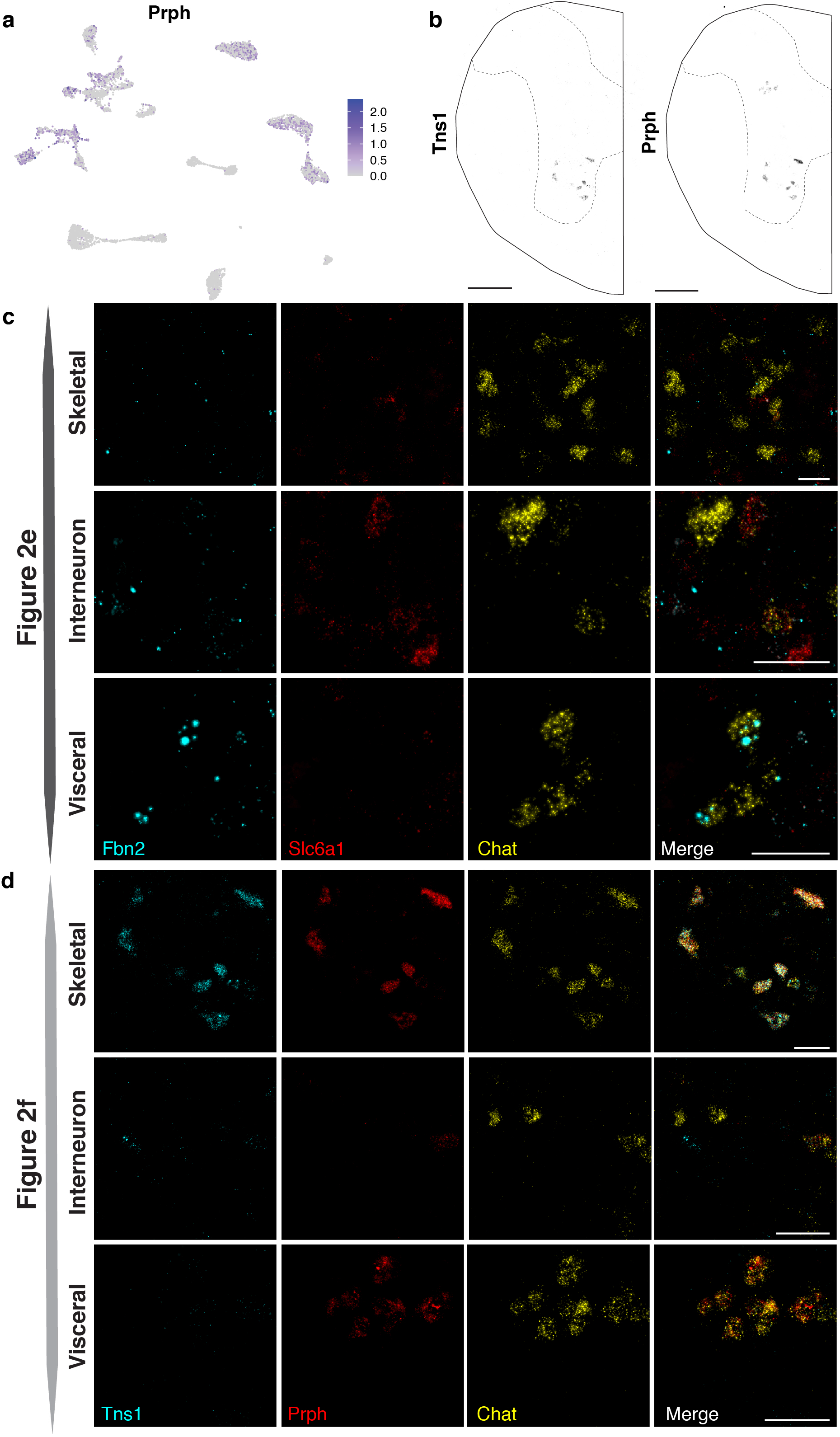
Expression of cholinergic subtype markers. **a**, A feature plot showing extent expression of *Prph* in our dataset across all cholinergic clusters. **b**, Low magnification images of *Tns1* expression, localized to the ventral horn, and *Prph* in ventral and lateral horn of thoracic spinal cord. **c**, High magnification images of skeletal MNs (ventral horn), interneurons (intermediate zone), and visceral MNs (lateral horn) from Figure 2e showing expression of *Fbn2* (cyan), *Slc6a1* (red), and *Chat* (yellow) in each channel individually, and merged. **d**, High magnification images of skeletal MNs (ventral horn), interneurons (intermediate zone), and visceral MNs (lateral horn) from Figure 2f showing expression of *Tns1* (cyan), *Prph* (red), and *Chat* (yellow) in each channel individually, and merged. Low magnification scale bars, 200 μm. High magnification scale bars, 50 μm.

**Extended Data Fig. 5.**
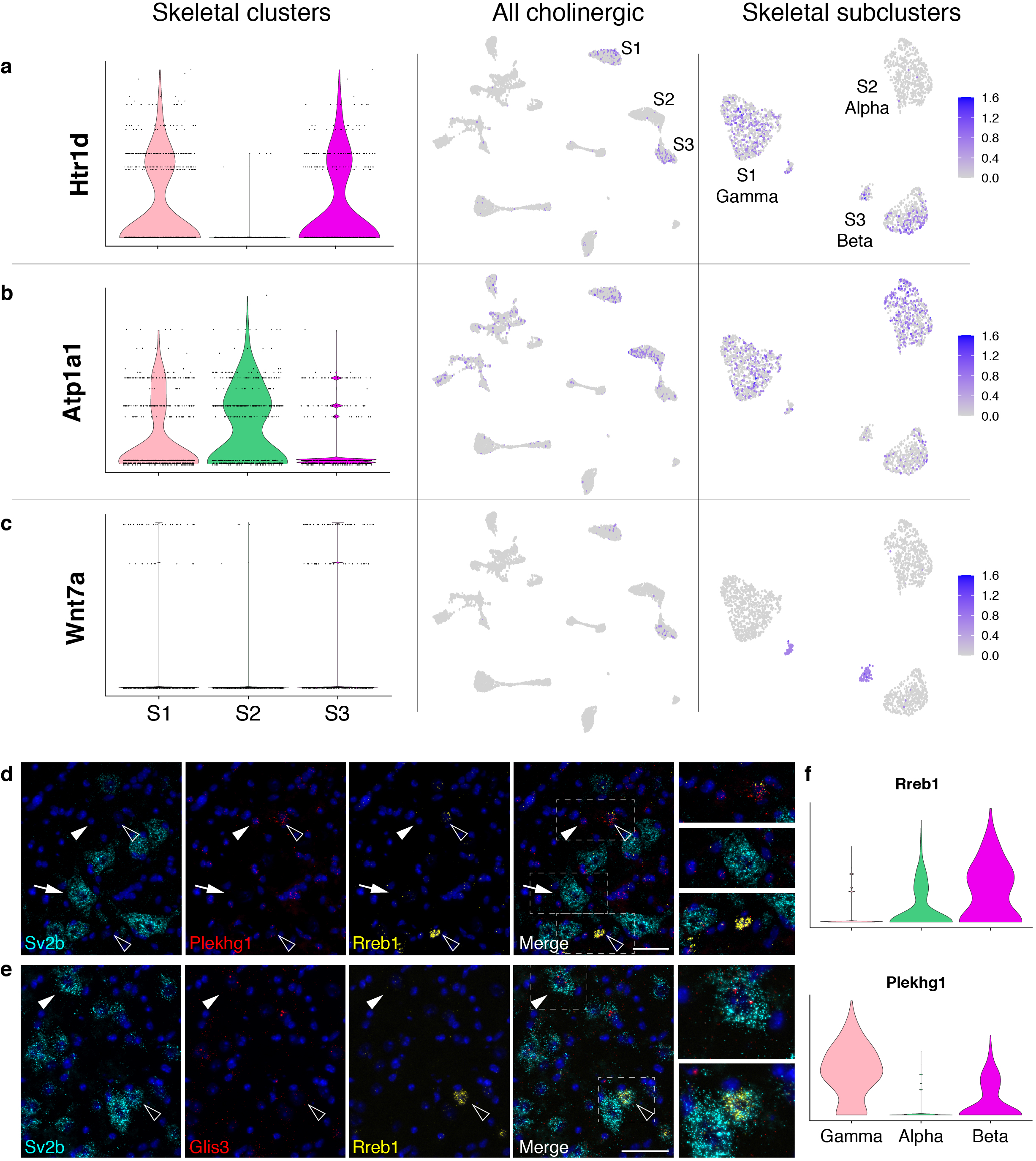
Expression of known and novel skeletal subtype markers in our dataset. **a**, The GDNF family receptor alpha 1 *(Gfra1)* has been described as a marker mainly expressed in gamma MNs, but is widely expressed in our nuclear sequencing dataset. **b**, The ATPase Na+/K+ transporting alpha 1 (*Atp1a1*) is considered a marker for alpha motor neurons, but is expressed across skeletal MNs in our dataset. **c**, *Wnt7a* is known to be expressed selectively in gamma MNs starting at late embryonic stages, but is found in small sub-clusters of both gamma and beta MNs in our dataset. **d**, Sacral spinal cord images show that *Glis3* (red) expression is restricted to a subset of alpha MNs (*Sv2b*+, cyan) (arrowhead, top). **e**, Lumbar spinal cord images show that *Sv2b* (cyan) labels alpha MNs (arrow, middle); *Rreb1* (yellow) alone or co-expressed with *Plekhg1* (red) represents beta MNs (open arrowhead, top); *Plekhg1* alone represents gamma MNs (arrowhead, top). A subset of alpha MNs express *Rreb1* (yellow) (open arrowhead, bottom). **f**, Violin plots of *Rreb1* and *Plekhg1* expression in alpha, beta, and gamma MNs. Scale bars, 50 μm.

**Extended Data Fig. 6.**
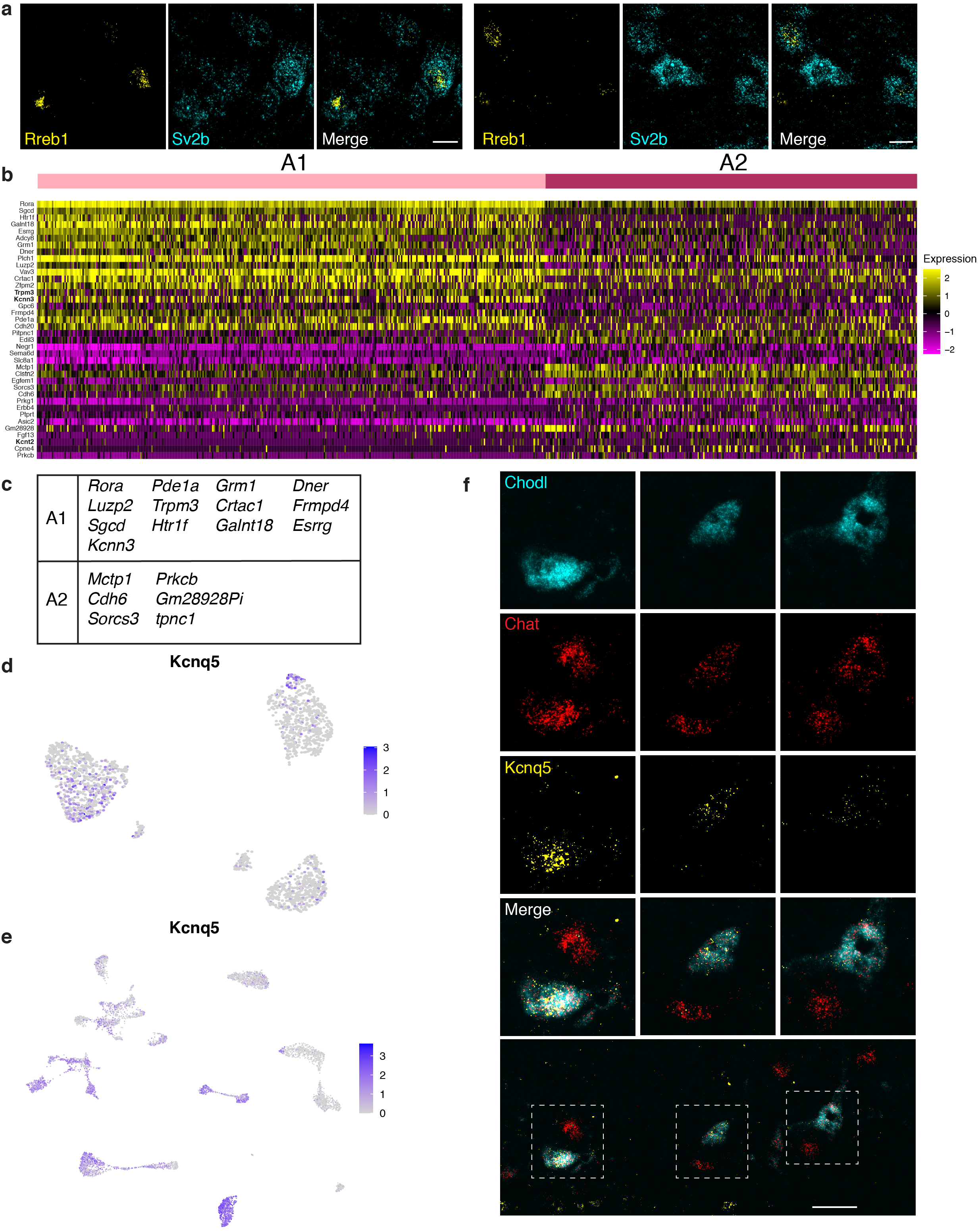
Skeletal clusters A1 and A2 express many shared novel markers as *Chodl*+ and *Rreb1*+ cells, and a candidate marker for fast alphas co-expresses with *Chodl* in ventral horn. **a**, Additional examples of slow-firing alpha MNs co-expressing *Rreb1* (yellow) and *Sv2b* (cyan) in the ventral horn of thoracic and lumbar spinal cord. Scale bars, 20 μm. **b**, Heat map showing expression of top genes differentiating between clusters A1 and A2. Genes in bold font are ion channels. **c**, Markers shared between this analysis of genes distinguishing A1 and A2, and the analysis in Fig. 4g of genes enriched in *Rreb1*+ vs *Chodl*+ cells. **d**, *Kcnq5* labels gamma MNs, but is highly expressed in a subset of cluster A2 alpha MNs. **e**, *Kcnq5* is widely expressed across cholinergic neurons. **f**, *Chodl* (cyan) and *Kcnq5* (yellow) co-express with a subset of large diameter *Chat*+ (red) neurons in the ventral horn of lumbar spinal cord. Other large diameter *Chat*+ neurons without *Chodl* or *Kcnq5* expression are likely other subtypes of skeletal MNs. Scale bar, 50 μm.

**Extended Data Fig. 7.**
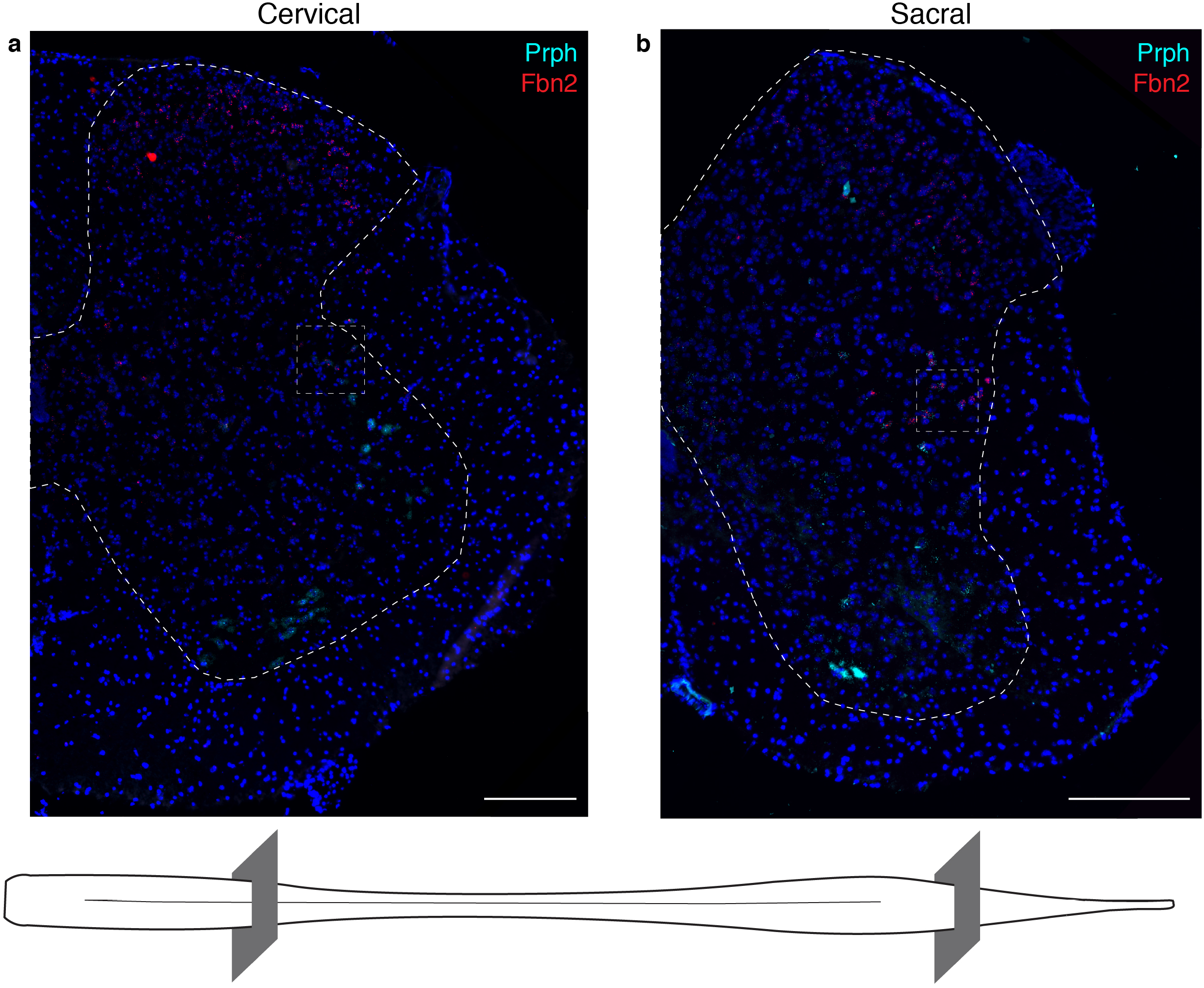
Localization of *Sst*+ and *Bnc2*+ visceral motor neurons. **a**, *Bnc2* co-expression with *Prph* and *Fbn2* was observed in the lateral horn in cervical spinal cord, but not in lower spinal cord levels. **b**, *Sst* co-expression with *Prph* and *Fbn2* was observed in the sacral spinal cord, but was not seen above lower lumbar levels. Low magnification images of Fig. 6e and 6d, respectively. The level of the spinal cord is evident by the shape of the section and its grey matter. Scale bars, 200 μm.

**Extended Data Fig. 8.**
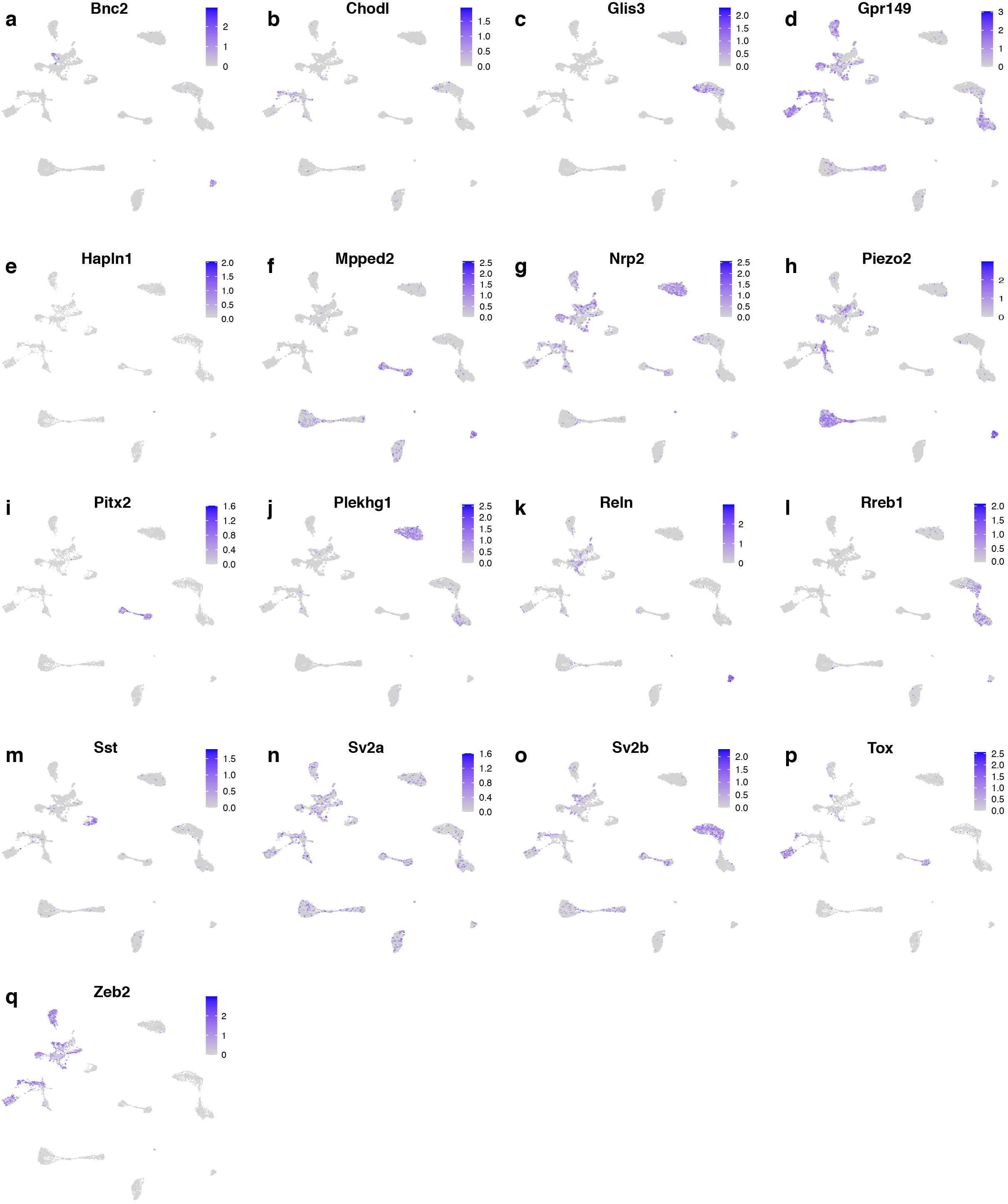
Nuclear expression across all cholinergic neurons sequenced for genes discussed in our analysis. **a-q**, UMAPs showing expression of genes discussed in our analysis across all cholinergic nuclei sequenced. Only displaying UMAPs not shown elsewhere. *Bnc2*, *Chodl*, *Glis3*, *Gpr149*, *Hapln1*, *Mpped2*, *Nrp2*, *Piezo2*, *Pitx2, Plekhg1*, *Reln*, *Rreb1*, *Sst, Sv2a*, *Sv2b*, *Tox*, *Zeb2*.

**Supplementary Table 1.** Summary of number of nuclei sequenced in this study.

| **All** |  | **All cholinergic** |  | **Skeletal** |  | **Visceral** |  |
| --- | --- | --- | --- | --- | --- | --- | --- |
| Cluster | Count | Cluster | Count | Cluster | Count | Cluster | Count |
| 0 | 1615 | I1 | 656 | A1 | 368 | 0 | 359 |
| 1 | 1527 | I2 | 616 | A2 | 270 | 1 | 329 |
| 2 | 924 | I3 | 303 | B1 | 378 | 2 | 306 |
| 3 | 842 | I4 | 233 | B2 | 104 | 3 | 278 |
| 4 | 838 | I5 | 175 | B3 | 74 | 4 | 228 |
| 5 | 792 | I6 | 83 | G1 | 411 | 5 | 224 |
| 6 | 755 | I7 | 176 | G2 | 390 | 6 | 219 |
| 7 | 683 | I8 | 21 | G3 | 41 | 7 | 208 |
| 8 | 650 | V1 | 467 |  |  | 8 | 151 |
| 9 | 631 | V2 | 457 |  |  | 9 | 114 |
| 10 | 615 | V3 | 454 |  |  | 10 | 91 |
| 11 | 566 | V4 | 328 |  |  | 11 | 71 |
| 12 | 540 | V5 | 279 |  |  | 12 | 40 |
| 13 | 459 | V6 | 232 |  |  | 13 | 24 |
| 14 | 336 | V7 | 220 |  |  |  |  |
| 15 | 324 | V8 | 205 |  |  |  |  |
| 16 | 303 | S1 | 841 |  |  |  |  |
| 17 | 277 | S2 | 631 |  |  |  |  |
| 18 | 236 | S3 | 564 |  |  |  |  |
| 19 | 230 |  |  |  |  |  |  |
| 20 | 227 |  |  |  |  |  |  |
| 21 | 222 |  |  |  |  |  |  |
| 22 | 205 |  |  |  |  |  |  |
| 23 | 176 |  |  |  |  |  |  |
| 24 | 172 |  |  |  |  |  |  |
| 25 | 161 |  |  |  |  |  |  |
| 26 | 137 |  |  |  |  |  |  |
| 27 | 134 |  |  |  |  |  |  |
| 28 | 89 |  |  |  |  |  |  |
| 29 | 52 |  |  |  |  |  |  |
| 30 | 20 |  |  |  |  |  |  |
| **TOTAL:** | **14738** |  | **6941** |  | **2036** |  | **2642** |

